# Intracellular aggregation of the transthyretin protein is limited by Hsp70 chaperones

**DOI:** 10.64898/2025.12.14.694244

**Authors:** Claire M. Radtke, Adam S. Knier, Sean A. Martin, Jane E. Dorweiler, Matt E. Hudson, Anita L. Manogaran

## Abstract

Extracellular amyloid deposits are a hallmark feature of systemic protein aggregation diseases such as transthyretin amyloidosis (ATTR). However, emerging evidence suggests that extracellular transthyretin (TTR) aggregates are internalized and result in an intracellular stress response, including elevated Hsp70 levels. While ATTR research has predominantly focused on extracellular TTR amyloid, our understanding of TTR aggregation inside the cell is poorly explored. To better understand how intracellular chaperones impact intracellular TTR, we used a yeast model that expresses TTR fused eGFP (TTR-eGFP) within the cytoplasm. Since the Hsp70 chaperone family, and co-chaperones the J-domain proteins (JDPs) and Hsp110, act as a disaggregase *in vitro,* we asked how these molecular chaperones impact TTR aggregation intracellularly *in vivo*. TTR-eGFP forms detergent-soluble high molecular weight (HMW) aggregates in yeast that have biochemistry profiles similar to human patient TTR. While knockdown of the JDP, Sis1, and deletion of the Hsp110, Sse1, appear to slightly increase TTR-eGFP aggregation, the loss of two major yeast Hsp70s, Ssa1 and Ssa2, lead to a significant increase in the size of HMW species. Taken together, our data suggest that Hsp70s limit the formation of HMW TTR aggregates in the intracellular environment. Based on our results, it is possible that the age-related decline of protein homeostasis, including Hsp70s, may promote the intracellular aggregation of TTR.

## Introduction

Extracellular aggregates are a hallmark of many age-related conditions such as neurodegenerative diseases like Alzheimer’s disease (AD) and systemic diseases like transthyretin amyloidosis (ATTR). However, research shows that the internalization of protein aggregates into cells may contribute to disease pathology. For example, the amyloid-β (Aβ) protein forms extracellular amyloid in AD but is also internalizes from the extracellular space via endocytosis and accumulates within neurons (LaFerla, Green et al. 2007, Hivare, Mujmer et al. 2023). This endocytic uptake can lead to lysosomal membrane permeabilization, ultimately releasing Aβ into the cytoplasm where it contributes to cellular dysfunction (Liu, Zhou et al. 2010, De Kimpe, van Haastert et al. 2013, Gallego Villarejo, Bachmann et al. 2022). Similarly, the transthyretin (TTR) protein, which normally circulates as a tetramer, but can dissociate into monomers and form extracellular amyloid. While the extracellular amyloid can infiltrate tissues, the amyloid has also been shown to accumulate intracellularly in patient tissues, including cardiac fibroblasts (Misumi, Ando et al. 2013) and Schwann cells (Goncalves, Costelha et al. 2014, Lu, Mak et al. 2024). Furthermore, the exposure of cultured neuroblastoma and glial cells to recombinant TTR oligomers is associated with cytotoxicity (Sousa, Cardoso et al. 2001, Sousa and Saraiva 2001, Andersson, Olofsson et al. 2002, Reixach, Deechongkit et al. 2004, Sorgjerd, Klingstedt et al. 2008, Dasari, Hughes et al. 2019), suggesting that TTR internalization could negatively impact the cell. Despite these findings, our understanding of TTR aggregation inside cells and how cells respond to TTR aggregation remains limited.

Cells rely on molecular chaperones for protein refolding, sequestration, disaggregation, and targeting proteins for degradation. A family of molecular chaperones called Hsp70s are neuroprotective against internally expressed Aβ in primary neuronal culture and transgenic mouse models (Magrane, Smith et al. 2004, Hoshino, Murao et al. 2011), suggesting that molecular chaperones like Hsp70s could mitigate disease-associated protein aggregation. Similarly, research shows that the molecular chaperone response may influence TTR aggregation. For example, ATTR patient biopsies show that tissues exposed to extracellular TTR deposits have increased levels of the master heat shock response transcription factor, Hsf1, which regulates the expression of many intracellular molecular chaperones (Santos, Magalhaes et al. 2008). ATTR mouse models in *hsf-1* null backgrounds also show extensive TTR aggregation (Santos, Fernandes et al. 2010). Furthermore, Hsp70 levels are elevated in cells surrounded by TTR deposits in ATTR patient biopsies (Santos, Magalhaes et al. 2008) and in induced pluripotent stem cells derived from ATTR patients (Leung, Nah et al. 2013), underscoring the impact that TTR has on the intracellular stress response. However, how these intracellular chaperone responses impact intracellular TTR aggregation is not well characterized.

Hsp70 is an ATP hydrolyzing chaperone best known for its ability to disaggregate a wide range of substrates *in vitro*, including luciferase, the yeast Sup35 protein, and several disease-associated fibrils (Shorter 2011, Gao, Carroni et al. 2015, Nachman, Wentink et al. 2020, Beton, Monistrol et al. 2022). Hsp70 disaggregation requires cooperation of J-domain protein (JDP) and Hsp110 co-chaperones. JDPs deliver substrates to Hsp70 and stimulate Hsp70’s ATPase activity (Cheetham and Caplan 1998, Fan, Lee et al. 2003, Kampinga and Craig 2010), while Hsp110 acts as a nucleotide exchange factor (NEF) that promotes ADP release, resetting Hsp70 for another round of substrate processing (Raviol, Sadlish et al. 2006, Andreasson, Fiaux et al. 2008). While Hsp70 has been shown to block Aβ and tau variant aggregation *in vitro* (Evans, Wisen et al. 2006, Kundel, De et al. 2018), studies in yeast show that loss of two major Hsp70s, Ssa1 and Ssa2, leads to the abnormal accumulation of model aggregating proteins and endogenous proteins (Andersson, Eisele-Burger et al. 2021, Rolli, Langridge et al. 2024, Buchholz, Martin et al. 2025). Despite Hsp70’s ability to mitigate protein aggregation, it is unclear if Hsp70 plays a role in limiting intracellular TTR aggregation.

To understand how Hsp70 and its co-chaperones manage TTR aggregation inside the cell, we turned to the yeast model. Yeast are advantageous as they are a simple eukaryotic system that exhibit rapid aggregation TTR. The expression of TTR fused to GFP (TTR-eGFP) in a wildtype yeast with unmodified Hsf1 levels results in visible detectable fluorescent puncta and biochemically detectable TTR aggregates in late log or stationary cultures (Verma, Girdhar et al. 2018, Knier, Davis et al. 2022). In contrast, mouse models exhibit long times for TTR aggregation, have ectopic aggregation, and often rely on *hsf-1* knockouts (Santos, Magalhaes et al. 2008, Santos, Fernandes et al. 2010, Goncalves, Costelha et al. 2014, Murakami, Ito et al. 2023). In this study, the majority of TTR-eGFP exists in high molecular weight (HMW) SDS-soluble species, and do not exhibit extensive SDS-resistance observed with other amyloid forming proteins. Using biochemical and microscopy approaches, depletion of the yeast JDP protein, Sis1, or deletion of the yeast Hsp110, Sse1, mildly increases the size of TTR-eGFP aggregates. However, loss of two Hsp70 proteins, Ssa1 and Ssa2, results in the presence of large HMW SDS-soluble TTR-eGFP species. Our data suggests that Hsp70s limit intracellular TTR aggregation in the cell, adding to the growing evidence that molecular chaperones play a role in limiting disease-associated protein aggregation.

## Results

### C-terminal eGFP tag enables strong TTR detection

In yeast, expression of the human TTR (amino acids 21–147) has been visualized by fluorescent microscopy using a C-terminal fluorescent eGFP tag (TTR-eGFP) (Derkatch, Uptain et al. 2004, Verma, Girdhar et al. 2018, Knier, Davis et al. 2022). Although the addition of an eGFP tag does not affect TTR’s ability to form tetramers or fibrils *in vitro* (Duan, Li et al. 2023), we wanted to compare the aggregation of TTR-eGFP to untagged TTR in yeast. Untagged TTR was expressed under a constitutive GPD promoter in the 74D-694 genetic background, which has been used to study aggregating proteins such as yeast prions (Chernoff, Lindquist et al. 1995). Wildtype strains transformed with untagged wildtype TTR (TTR^WT^) had similar growth compared to the non-toxic and well characterized TTR-eGFP (Verma, Girdhar et al. 2018, Knier, Davis et al. 2022) (Supplemental Fig. 1A). Additionally, TTR familial mutations known to enhance aggregation (Dasari, Hung et al. 2019) and another mutation L110Q (Knier, Davis et al. 2022) also showed similar growth to TTR-eGFP. Western blot analysis shows a band at the expected size of monomeric TTR-eGFP (∼41 kDa) using a polyclonal TTR antibody specific to the whole TTR protein. In contrast, monomeric untagged TTR^WT^ (∼14 kDa) gave a much weaker signal (Supplemental Fig. 1B). Some aggregation-prone proteins require strong denaturants to expose epitopes for antibody detection. Therefore, we compared the detection of untagged TTR^WT^ from lysates prepared under non-denaturing conditions and in the presence of a strong denaturant, 8M urea. However, urea treatment failed to produce a detectable signal for untagged TTR^WT^, and untagged TTR mutants (Supplemental Fig. 1C). Altogether, untagged TTR is poorly detected, and therefore we continued our analyses with TTR-eGFP.

### TTR-eGFP forms high molecular weight (HMW) SDS-soluble species

The treatment of yeast lysates with SDS has been reported to yield monomeric and SDS-resistant TTR-eGFP (Verma, Girdhar et al. 2018, Knier, Davis et al. 2022). However, no study has evaluated the TTR-eGFP aggregation state in yeast under native conditions. Therefore, native polyacrylamide gel electrophoresis (native PAGE) was used to separate proteins under non-denaturing conditions. In humans, TTR circulates as a tetramer and is rarely detected as a monomer. Similarly in untreated yeast lysates, no signal equivalent to monomeric TTR-eGFP (approximately 41 kDa) was detected (Fig. 1A). Instead, several bands are detected as well as a smear starting near 242 kDa and extending to the top of the gel. Interestingly, a recent study analyzing ATTR patient plasma with native PAGE showed that TTR exists as a similar smear (called oligomers) and at the top of gel (called high molecular weight, or HMW TTR). Both of these features were absent in control patients (Pedretti, Wang et al. 2024). The similarity between the patient TTR and our TTR-eGFP native PAGE profiles suggests that the TTR-eGFP yeast resembles disease-specific TTR aggregation seen in ATTR patients. Therefore, we refer to the TTR-eGFP smear as oligomers and TTR at the top of the gel at HMW TTR, although we recognize these may not be homogenous TTR species.

**Figure 1.**
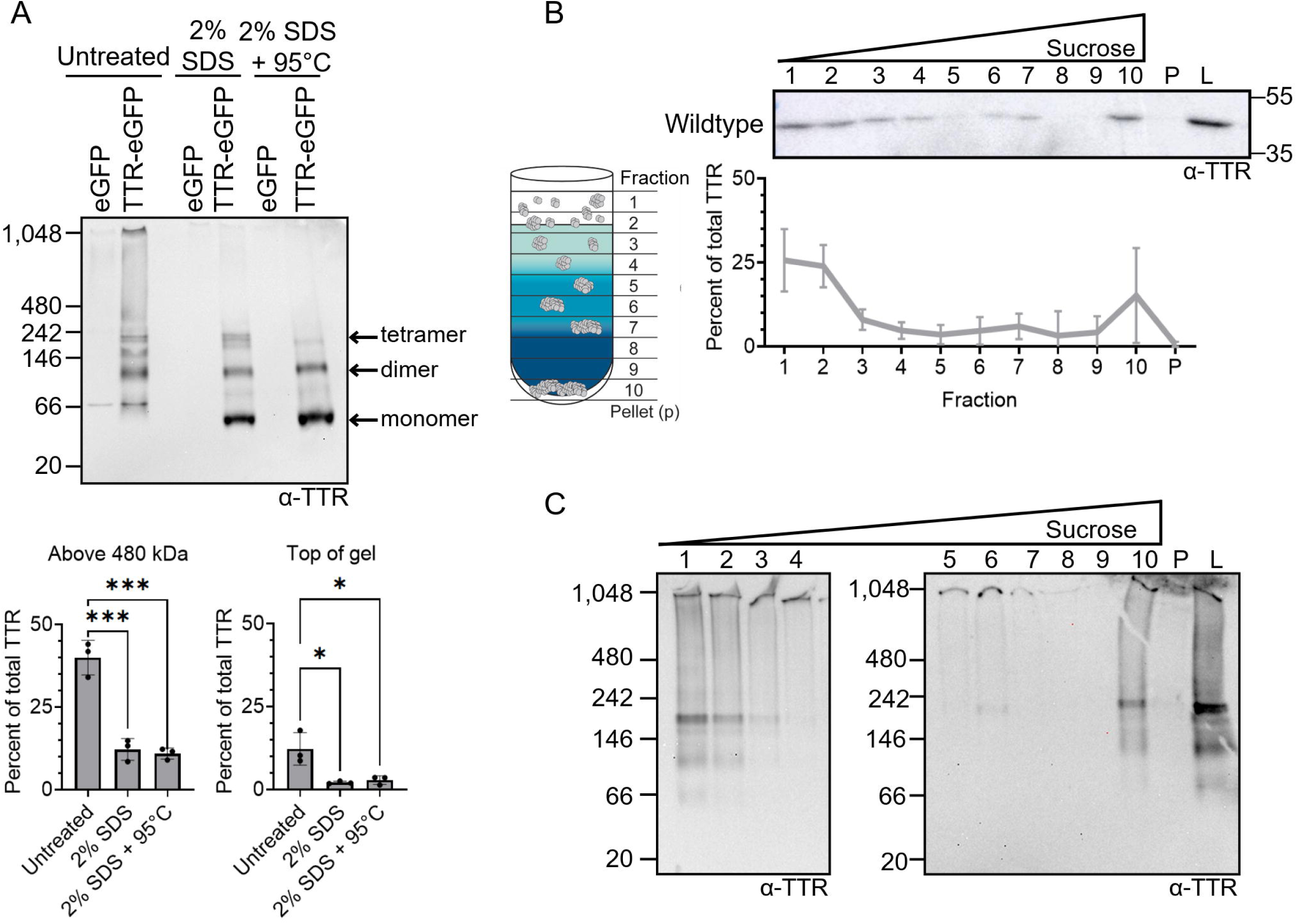
TTR-eGFP forms high molecular weight (HMW) SDS-soluble species. A) Wildtype strains were transformed with eGFP (p3141) or TTR-eGFP. Lysates were analyzed by native PAGE under the following conditions: untreated, treated with 2% SDS, or 2% SDS incubated at 95°C for 8 minutes. Samples were resolved on 4–20% tris-glycine native PAGE and immunoblotted with polyclonal anti-TTR antibody. Estimated monomeric, dimeric, and tetrameric TTR-eGFP sizes are labeled. *Bottom:* Quantification was performed by measuring the intensity of TTR signal between 480 kDa and the top of the gel (Above 480 kDa) or the area around the lane wells (Top of gel) and normalizing it to the total TTR signal in each lane. Dots represent independent trials and shown as means ± SD. Statistical analysis was performed with a one-way ANOVA and Tukey post hoc test (* p < 0.05, *** p < 0.001). Unlabeled comparisons are not significant. B) *Left:* Schematic of protein sedimentation in discontinuous sucrose gradients composed of 10%, 40%, and 60% sucrose. Lysate is loaded on top of the gradient, and proteins separate by density during centrifugation. Equal-volume fractions are collected after sedimentation. *Right:* Individual fractions (1–10), pelleted protein (P), and whole-cell lysates (L) were analyzed by SDS-PAGE and immunoblotted with monoclonal anti-TTR antibody. Shown is a representative image from seven independent biological trials. The TTR intensity from each fraction was normalized to the combined total TTR signal in fractions 1-10 and pellet and mean ± SD is graphed. C) Lysates from wildtype strains transformed with TTR-eGFP were subjected to sucrose gradient sedimentation, and fractions were analyzed by 4–20% tris-glycine native PAGE and immunoblotted with a polyclonal anti-TTR antibody. Shown are representative images from three independent biological trials.

Since previous data has reported that TTR-eGFP is also SDS-soluble (Knier, Davis et al. 2022), we next tested whether any TTR-eGFP signal is lost on native PAGE upon treatment with SDS. Indeed, 2% SDS treatment resulted in a loss of signal from the oligomeric-like smear and HMW TTR-eGFP, and the appearance of a strong band at the expected size of monomeric TTR-eGFP (approximately 41 kDa; Fig. 1A). To quantify the SDS-soluble TTR-eGFP, we selected the area above the 480 kDa marker because the 242 kDa marker migrated too close to an SDS-resistant band. Using the 480 kDa marker, we observed approximately 40% of the TTR-eGFP signal migrates above 480kDa in non-treated lysates (Fig. 1A). The addition of SDS showed a significant threefold reduction in any TTR signal above 480 kDa. Specifically, the HMW TTR-eGFP signal at the top of the gel, likely representing protein too large to sufficiently enter the gel, exhibited a significant twofold decrease in signal with SDS (Fig. 1A). These data suggest that oligomeric-like and HMW TTR-eGFP are SDS-soluble.

In addition to the strong monomeric signal, we also observed three prominent bands with SDS treatment. A doublet appeared with sizes between ∼160-200 kDa. We estimate the lower doublet band to be the tetramer, which is consistent with previous reports of a detergent resistant TTR tetramer (Henze, Homann et al. 2015). The upper doublet band may represent a post-translationally modified tetramer, such as previously described cysteine modifications (Lim, Prokaeva et al. 2002, Kingsbury, Laue et al. 2008). Additionally, the 90 kDa SDS-resistant band is estimated to be a dimer (Fig. 1A), which is similar to observations from recombinant and patient samples (Gao, Xie et al. 2022, Jayaweera, Sahin et al. 2025). Further incubation of SDS-treated samples at 95°C resulted in the loss of the tetramer-like bands, and the persistence of the dimer-like band, indicating that the tetramer-like band is heat sensitive. Lastly, we tested whether a native PAGE system optimized for resolving HMW protein (tris-acetate native PAGE) could improve the resolution of the smear and HMW TTR-eGFP. Similar to our initial native PAGE analysis, this approach revealed a continuous oligomer-like smear of TTR-eGFP extending up to the top of the gel (Supplemental Fig. 2A-B). Anti-GFP blots confirmed that these smears originate from TTR-eGFP, whereas samples expressing eGFP alone appeared as a single band.

The presence of TTR-eGFP at the top of the gel in both native PAGE systems strongly suggests that there are HMW TTR-eGFP species that are above the detection limit of native PAGE. Therefore, discontinuous sucrose gradients were used to separate the HMW TTR-eGFP that are too large to enter native PAGE, as sucrose gradients have been shown to trap HMW species in the interfaces between different sucrose concentrations (Fig. 1B) (Dorweiler, Oddo et al. 2020). Here, we find that approximately 50% of the total TTR-eGFP is present in fraction 1 and 2, with the remainder of TTR-eGFP able to penetrate into the gradient (Fig. 1B), similar to our previous data (Knier, Davis et al. 2022). While there is a relatively low percent of total TTR-eGFP distributed among fractions 3-9, approximately 18% of total TTR-eGFP is detected in fraction 10. To better understand the distribution of molecular weights found in each fraction, all sucrose gradient fractions were subjected to native PAGE analysis (Fig. 1C). Fraction 1 and 2 on native PAGE had TTR-eGFP banding patterns that resembled unfractionated whole lysates (Fig. 1A), including dimer-like and tetramer-like bands, and oligomer-like smears extending to the top of gel. In contrast, fractions 3–9 predominantly showed HMW TTR-eGFP almost exclusively at the top of gel. The presence of HMW TTR-eGFP in fractions 3-9 suggests that TTR-eGFP species exceeds the detection limit of native PAGE, and result in the distribution throughout the sucrose gradient. Interestingly, fraction 10 exhibited a TTR-eGFP pattern similar to the unfractionated whole lysate. We suspect that because sucrose gradients are detergent free techniques, fraction 10 may represent TTR-eGFP associated with other proteins and lipids. Indeed, treatment with 0.1% Triton X-100, a non-ionic detergent that disrupts membranes and solubilizes protein-lipid interactions, dramatically, but not significantly, reduced TTR-eGFP in fraction 10 compared to other fractions (Supplemental Fig. 2C). Therefore, we suspect that TTR-eGFP in Fraction 10 likely reflects the disruption of TTR and detergent labile interactions.

While previous work has suggested that TTR-eGFP exists in both SDS-resistant and SDS-soluble species (Verma, Girdhar et al. 2018, Knier, Davis et al. 2022), our native PAGE and sedimentation analysis provides more detailed information about these species. Our analysis suggests that only the tetramer-like and dimer-like TTR-eGFP species are SDS-resistant. On the contrary, TTR-eGFP exists in SDS-soluble oligomer-like and HMW species that are above 480kDa in size, as well as TTR-eGFP possibly associated with lipid-like or Triton X-100 soluble entities. Altogether, TTR-eGFP appears to form SDS-soluble aggregate species that can be detected across both native PAGE and sucrose gradients.

### Knockdown of the yeast J-domain protein, Sis1, slightly shifts TTR sedimentation into the gradient

Having established several techniques to analyze TTR-eGFP aggregate species, we next asked whether molecular chaperones influence TTR-eGFP aggregation in yeast. We first focused on the essential yeast cytosolic JDP, Sis1, as previous work suggests that Sis1 enables more efficient Hsp70-mediated disaggregation of model aggregating proteins in comparison to the other cytosolic JDP, Ydj1 (Shorter 2011, Wyszkowski, Janta et al. 2021). Since Sis1 is an essential gene, any strain lacking *SIS1* must be maintained with an exogenous *SIS1* copy. A 74D-694 *sis1Δ* deletion strain with a plasmid expressing a tetracycline-repressible *SIS1* (*sis1*-*Δ* [*TETrSIS1*]) was used for these studies (Aron, Higurashi et al. 2007). In the absence of doxycycline (doxy), *sis1*-*Δ* [*TETrSIS1*] showed an ∼1.8-fold increase in Sis1 steady-state levels compared to wildtype strains, yet TTR-eGFP levels appeared similar (Supplemental Fig. 3A). We suspect that the elevated Sis1 levels in *sis1*-*Δ* [*TETrSIS1*] likely reflects the constitutive expression from the tetracycline promoter in contrast to the stress-responsive endogenous Sis1 promoter in wildtype cells. And as expected, the addition of 10 μg/mL doxy to repress Sis1 transcription reduced Sis1 levels ∼17-fold without affecting TTR-eGFP steady state levels (Supplemental Fig. 3B).

To understand how Sis1 knockdown impacts TTR-eGFP aggregation, we utilized three different techniques: native PAGE, discontinuous sucrose gradients, and fluorescent microscopy. Native PAGE analysis between no doxy and doxy treated samples showed similar TTR-eGFP banding patterns, including the oligomer-like smear above 480 kDa and HMW TTR-eGFP at the top of the gel (Fig. 2A). No difference in TTR-eGFP migration was also observed by native PAGE in the presence of SDS treatment (Supplemental Fig. 4A). However, sucrose gradient analysis revealed a difference between untreated and doxy treated samples (Fig. 2B). In doxy treated samples, TTR-eGFP in fraction 1 decreased approximately threefold compared to untreated samples. Additionally, the loss of TTR-eGFP in fraction 1 was accompanied by a twofold increase in fraction 7 and fivefold increase in fraction 8 in doxy treated samples. The increase of TTR-eGFP at fraction 7 and 8 suggests that Sis1 knockdown leads to more TTR-eGFP trapped at the 40% and 60% sucrose solution interface. Lastly, to assess whether the addition of doxy induced changes in TTR-eGFP species beyond the detection limits of native PAGE and sucrose gradients, we used confocal microscopy to detect visible large fluorescent foci. Surprisingly, we find that even in the absence of doxy, the percentage of cells with TTR-eGFP foci was about 30% in the untreated *sis1*-*Δ* [*TETrSIS1*] strain, compared to the 4% of cells in wildtype (Fig. 2C). If the increase of cells with TTR-eGFP foci was solely due to increased Sis1 levels in the *sis1*-*Δ* [*TETrSIS1*] strain, then we would expect Sis1 knockdown to reduce the percent of cells with foci. However, the doxy induced knockdown of Sis1 resulted in no change in the percent of cells with foci, compared to untreated samples (Fig. 2C). Therefore, we suspect that there may be strain-specific differences between the *sis1*-*Δ* [*TETrSIS1*] and wildtype responsible for the differences in cells with visible foci. Overall, our data indicates that when comparing untreated to doxy treated samples, the Sis1 knockdown enhances TTR-eGFP penetration into the sucrose gradient, suggesting that a small portion of TTR exists in larger aggregates when Sis1 is not present.

**Figure 2.**
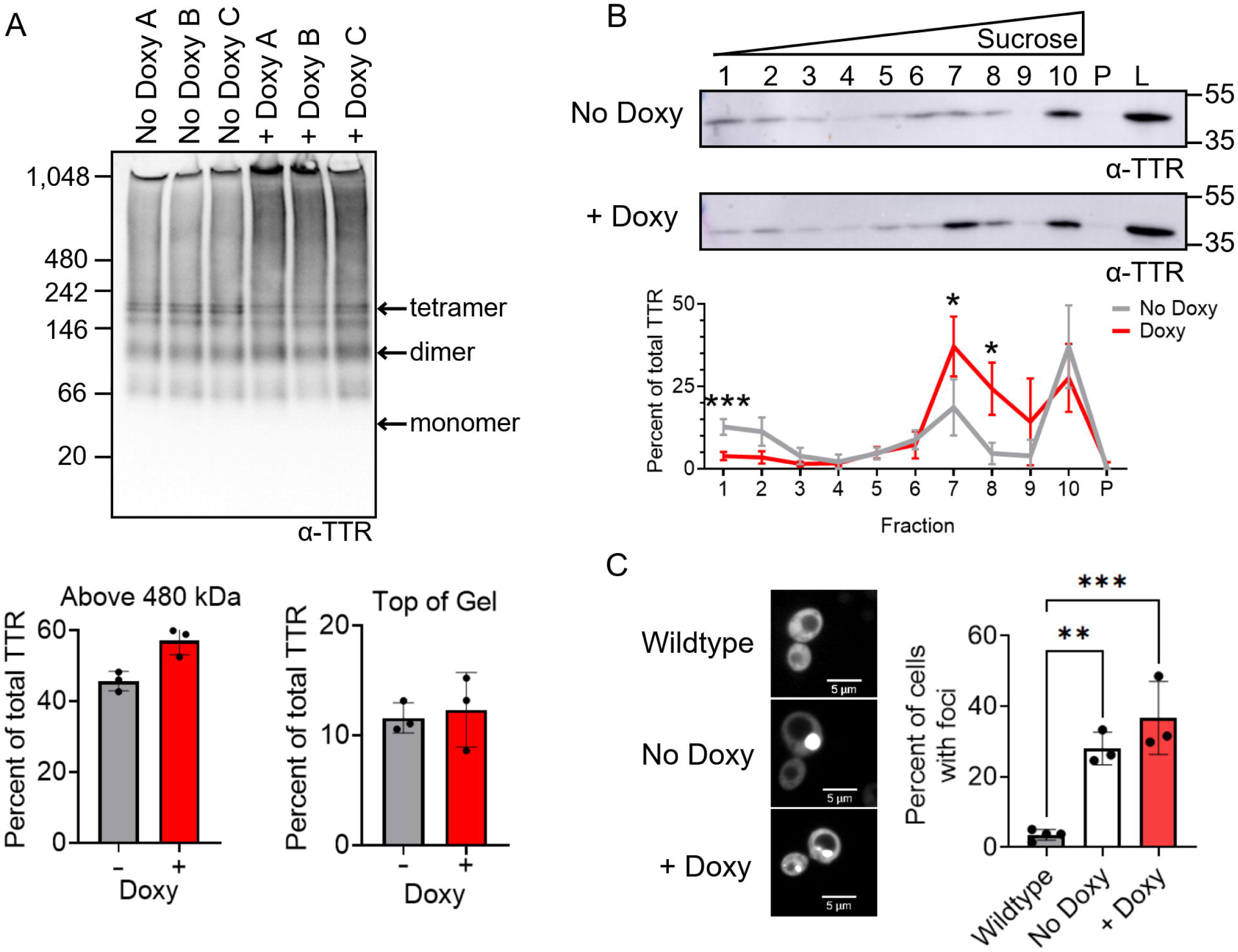
Sis1 knockdown results in loss of TTR-eGFP signal in fraction 1 and a gain in fractions 7 and 8. Paired *sis1*-*Δ* [*TETrSIS1*] cultures expressing TTR-eGFP were split, and either left untreated or treated with 10 µg/mL doxy overnight. A) Lysates were run on a 4–20% tris-glycine Native PAGE and immunoblotted in polyclonal anti-TTR antibody. Estimated monomeric, dimeric, and tetrameric TTR-eGFP sizes are labeled. Quantification was performed by measuring the area of the TTR signal of interest and normalizing to the total TTR signal in each lane. Dots indicate individual trials and shown as means ± SD. Statistical analysis was performed with a paired t-test (*p< 0.05). B) Untreated and 10 µg/mL doxy treated lysates were subjected to discontinuous sucrose gradient sedimentation. Fractions were loaded on SDS-PAGE and immunoblotted with a monoclonal anti-TTR antibody. Shown is a representative blot from six independent trials. The TTR signal from each fraction was normalized to the combined TTR signal in fractions 1-10 and pellet and graphed as mean ± SD. Statistical analysis was performed with a two-way repeated measures ANOVA, Geisser-Greenhouse correction, and a Sidak multiple comparisons test (* p < 0.05). C) *Left.* Indicated strains were imaged using spinning disk confocal microscopy at 630X magnification. Scale bars are shown. *Right.* Quantification of cells that contain TTR-eGFP foci, with each dot representing the mean from approximately 200 cells per trial. Graphs are drawn as mean ± SD. Statistical analysis were performed with a paired t-test (* p< 0.05) between doxy and non-doxy treated samples. All unlabeled comparisons are not significant. One-way ANOVA was used to compare doxy and non-doxy treated samples to wildtype strains, with Dunnett multiple comparison test.

### Deletion of the yeast Hsp110, Sse1, results in loss of TTR-eGFP in sucrose gradient fraction 1

In yeast, there are two major Hsp110 proteins, Sse1 and Sse2. While both proteins have similar sequences, Sse1 is constitutively expressed and Sse2 is stress responsive (Shaner, Trott et al. 2004, Raviol, Sadlish et al. 2006). However, the deletion of both Sse1 and Sse2 genes is lethal (Shaner, Wegele et al. 2005). Since the deletion of Sse2 did not change TTR-eGFP sucrose gradient sedimentation compared to wildtype strains (Supplemental Fig. 5), we assessed TTR-eGFP aggregation in *sse1Δ* strains. Immunoblotting with an Sse antibody, which detects both Sse1 and Sse2, resulted in a dramatic decrease in signal, which is consistent with a loss of Sse1, but not Sse2 (Ho, Baryshnikova et al. 2018) (Supplemental Fig. 6A). Additionally, steady state levels of TTR-eGFP, the endogenous yeast chaperone Hsp104, and the Ssa family were similar between in *sse1Δ* and wildtype cells (Supplemental Fig. 6A-B). In contrast, Sis1 levels were approximately twofold higher in *sse1Δ* strains with TTR-eGFP than in wildtype (Supplemental Fig. 6B), which suggests that the presence of TTR-eGFP may increase Sis1 steady state levels.

Native PAGE analysis of *sse1Δ* showed no change in TTR-eGFP banding patterns compared to wildtype, confirmed by quantification of total TTR-eGFP above 480 kDa and HMW at the top of the gel (Fig. 3A). Additionally, no difference in TTR-eGFP migration was detected by native page in the presence of SDS treatment between wildtype and *sse1Δ* (Supplemental Fig. 7A). Interestingly, sucrose gradient analysis of *sse1Δ* showed an eightfold loss of TTR-eGFP in fraction 1 and a roughly threefold increase in fraction 7 (Fig. 3B). However, there was no difference in the percent of cells with TTR-eGFP foci between wildtype and *sse1Δ* strains (Fig. 3C). As expected, cells expressing eGFP alone in either background exhibited almost no foci. Taken together, sucrose gradients revealed an overall loss of TTR-eGFP in fraction 1 and a gain in fraction 7 in *sse1Δ*.

**Figure 3.**
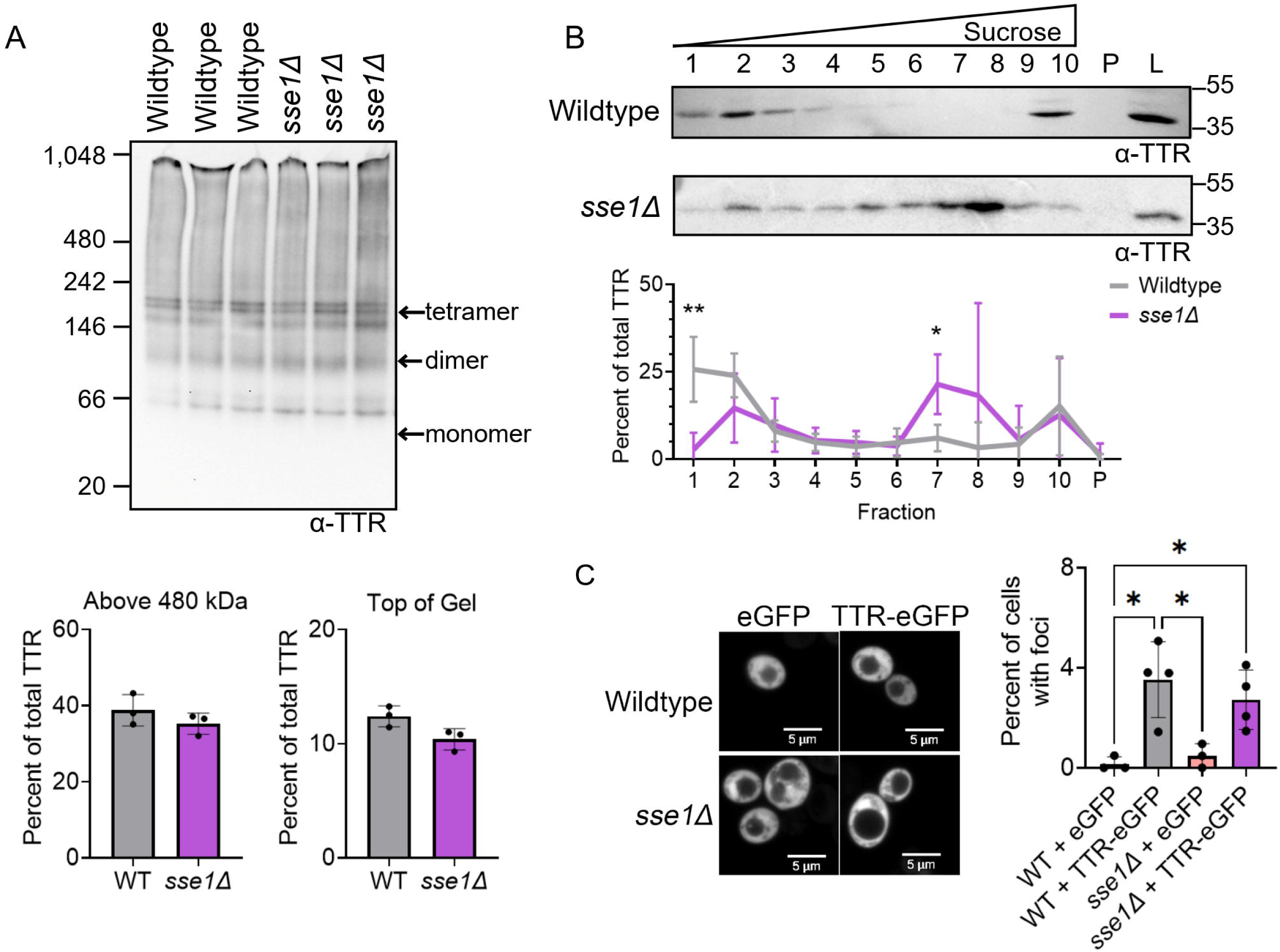
TTR-eGFP levels are reduced in fraction 1 and increased in fraction 7 in *sse1Δ*. Wildtype and *sse1Δ* strains were transformed with TTR-eGFP. A) Lysates were subjected to 4–20% tris-glycine Native PAGE and immunoblotted in polyclonal anti-TTR antibody. Quantification was performed by measuring the area of the TTR signal of interest and normalizing it to the total TTR signal in each lane. Dots indicate independent trials graphed as mean ± SD. Statistical analysis was performed with a t-test (*p< 0.05). B) Sucrose gradient sedimentation fractions were loaded on SDS-PAGE and immunoblotted with a monoclonal anti-TTR antibody. Shown is a representative blot from six independent trials. The TTR signal from each fraction was normalized to the combined TTR signal in fractions 1-10 and pellet and graphed as mean ± SD. Statistical analyses were performed with a two-way repeated measures ANOVA, Geisser-Greenhouse correction, and a Sidak multiple comparisons test (* p < 0.05). C) Indicated strains were imaged using spinning disk confocal microscopy at 630X magnification. Scale bars are shown. Quantification of cells containing TTR-eGFP foci, with each dot representing the mean from 200 cells per trial. Graphs are drawn as mean ± SD. Statistical analysis was performed with a one-way ANOVA and Tukey post hoc test (* p < 0.05). Unlabeled comparisons are not significant.

Given that Sse1 is well established as a nucleotide exchange factor (NEF) for Hsp70, we next asked whether the changes in TTR-eGFP sucrose gradient sedimentation in *sse1Δ* strains were due to Sse1 NEF activity. We utilized the Sse1 G233D mutation (Sse1^G223D^), which reduces both ATP (Shaner, Trott et al. 2004) and Ssa1 binding (Shaner, Wegele et al. 2005), thereby reducing Sse1 NEF activity. Both Sse1^G233D^ and the wildtype Sse1 (Sse1^WT^) control were expressed from plasmids under control of the endogenous Sse1 promoter, resulting in steady state levels comparable to the wildtype strain (Supplemental Fig. 7B). Similar to *sse1Δ* alone, expression of the empty vector (EV) in the *sse1Δ* strain resulted in very low TTR-eGFP detection in fraction 1 (Supplemental Fig. 8B). However, unlike *sse1Δ* alone, which showed 21% of total TTR-eGFP in fraction 7, only 3% of total TTR-eGFP was detected in fraction 7 when EV was expressed. We suspect that these differences in TTR-eGFP detected in fraction 7 may be due to the use of double-selective media or unknown effects associated with the presence of the two plasmids. The expression of Sse1^WT^ and Sse1^G233D^ increased the detection of TTR-eGFP in fraction 1 by 7-fold and ∼9-fold increase, respectively, compared to *sse1Δ* + EV (Supplemental Fig. 8C-D). Therefore, we suspect that Sse1-dependent loss of TTR-eGFP in fraction 1 is not dependent upon activities associated with ATP- and Hsp70- binding.

### Yeast Hsp70s, Ssa1 and Ssa2, influence TTR-eGFP aggregation

Ssa1 and Ssa2 are constitutively expressed cytoplasmic yeast Hsp70s (Werner-Washburne, Stone et al. 1987), which contribute to a range of protein quality control functions (Nollen and Morimoto 2002, Lee do, Sherman et al. 2016, Krakowiak, Zheng et al. 2018). The single deletion of Ssa1 resulted in no change in TTR-eGFP sucrose gradient sedimentation (Supplemental Fig.9A). Given that Ssa1 and Ssa2 have high sequence similarity and functional redundancy (Werner-Washburne, Stone et al. 1987), we next analyzed the double deletion of Ssa1 and Ssa2. The *ssa1Δssa2Δ* strain has been reported to have a slow growth phenotype, increased replicative aging (Andersson, Eisele-Burger et al. 2021), and increased protein aggregation of model aggregation and endogenous stress-responsive proteins (Andersson, Eisele-Burger et al. 2021, Rolli, Langridge et al. 2024, Buchholz, Martin et al. 2025). We found that in *ssa1Δssa2Δ*, Hsp104 levels almost doubled compared to wildtype (Supplemental Fig. 6A), consistent with reports of elevated heat shock response in *ssa1Δssa2Δ* (Matsumoto, Akama et al. 2005, Buchholz, Martin et al. 2025). Additionally, while Sse1/Sse2 and Sis1 steady state were similar between *ssa1Δssa2Δ* and wildtype strains, the TTR signal was reduced about ∼10-fold (Supplemental Fig. 6A, C). This finding is consistent with previous reports showing that the abundance of several endogenous yeast proteins, such as Pab1 and Sup35, are similarly decreased in *ssa1Δssa2Δ* (Buchholz, Martin et al. 2025), suggesting a general reduction in protein levels rather than a TTR-specific effect.

Native PAGE analysis showed that the percent of total TTR-eGFP protein above 480 kDa was similar between *ssa1Δssa2Δ* and wildtype strains. However, the percent of HMW TTR-eGFP at the top of the gel was modestly yet statistically higher in *ssa1Δssa2Δ*, increasing from 10% in wildtype compared to 14% in the *ssa1Δssa2Δ* (Fig. 4A). No difference in TTR-eGFP migration was detected by native PAGE in the presence of SDS treatment (Supplemental Fig. 9B). Sucrose gradient analysis revealed a significant reduction of TTR-eGFP in across fractions 1 through 4 in *ssa1Δssa2Δ* <u>strains</u> (Fig. 4B). This loss of TTR-eGFP in these fractions was accompanied by an increase in TTR-eGFP levels in fraction 7 and 10. Lastly, the percentage of cells containing TTR-eGFP foci showed a 3.75-fold increase in *ssa1Δssa2Δ* compared to wildtype strains (Fig. 4C).

**Figure 4.**
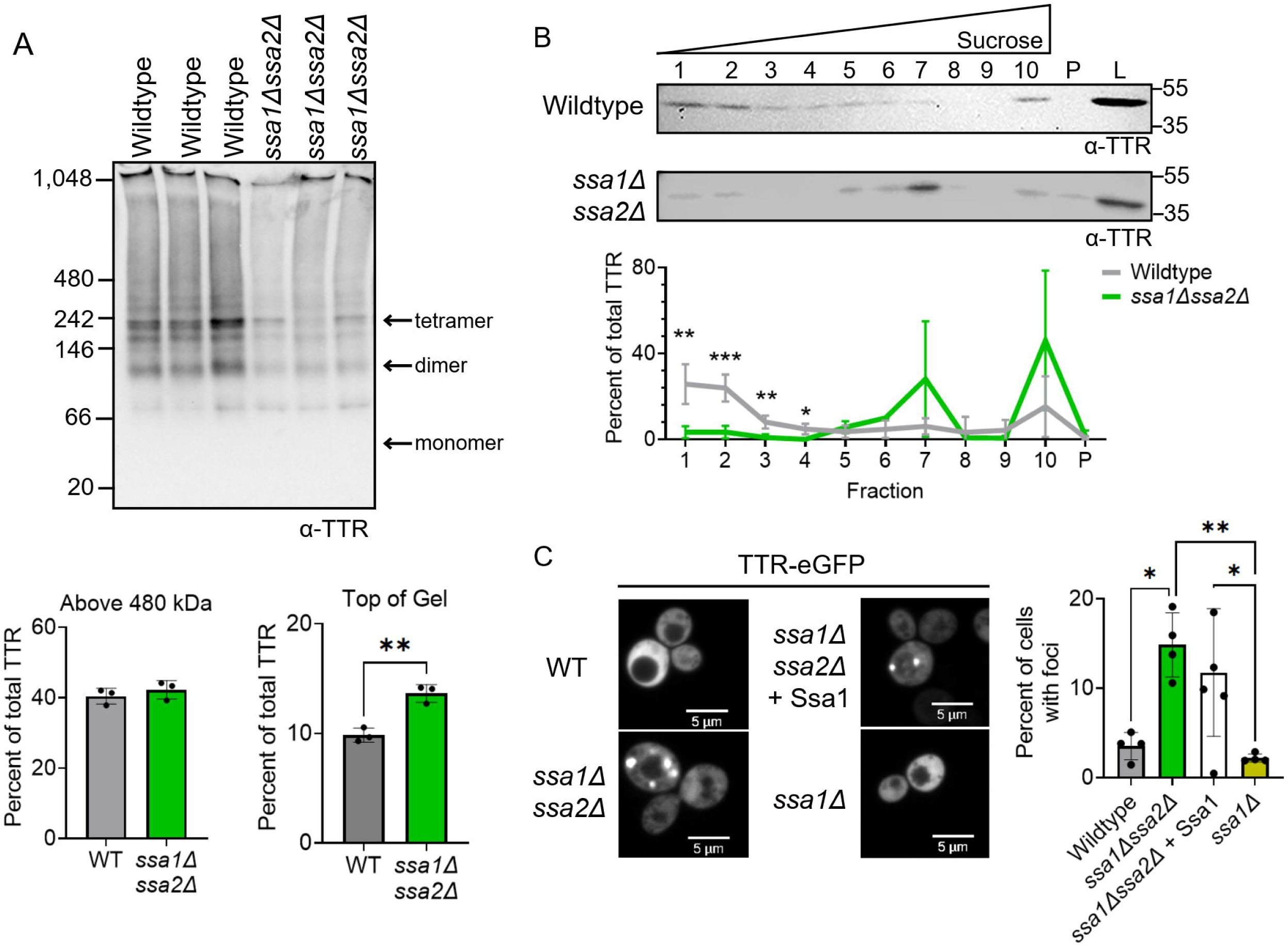
Larger TTR-eGFP species detected in *ssa1Δssa2Δ* strains. A-B) Lysates from wildtype and *ssa1****Δ****ssa2****Δ*** cultures expressing TTR-eGFP were analyzed by 4–20% tris-glycine native PAGE and immunoblotted with polyclonal anti-TTR antibody. Estimated monomeric, dimeric, and tetrameric TTR-eGFP sizes are labeled. Quantification was performed by measuring the area of the TTR signal of interest and normalizing it to the total TTR signal in each lane. Dots indicate independent trials and graphed as mean ± SD. Statistical analysis was performed with a t-test (**p<0.01). B) Lysates were subjected to discontinuous sucrose gradient sedimentation and fractions subjected to Western blot analysis. Shown is a representative blot from three independent trials. The TTR signal from each fraction was normalized to the combined TTR signal in fractions 1-10 and pellet and graphed as mean ± SD. Statistical analyses were performed with a two-way repeated measures ANOVA, Geisser-Greenhouse correction, and a Sidak multiple comparisons test (* p < 0.05). C) Cells were imaged using spinning disk confocal microscopy with a 63X objective. Scale bars are shown. Shown are representative images from at least three independent experiments. Dots represent the mean from independent trials of n=200 cells per trial. Graphs are drawn as mean ± SD. Statistical analysis was performed with a one-way ANOVA and Tukey post hoc test (* p < 0.05, ** p < 0.01). Unlabeled comparisons are not significant.

We next asked whether restoring Ssa1 expression in *ssa1Δssa2Δ* cells changes the percent of cells with TTR-eGFP foci. The introduction of a constitutively expressed Ssa1 into *ssa1Δssa2Δ* strains essentially is a strain that lacks Ssa2 expression. The *ssa1Δssa2Δ* +Ssa1 strain does not change the percent of cells with TTR-eGFP foci (Fig. 4C). This finding contrasts with previous results in which Pab1 foci in *ssa1Δssa2Δ* are rescued by Ssa1 overexpression (Buchholz, Martin et al. 2025). Analysis of the single deletion of Ssa1 (*ssa1Δ*) showed significantly less TTR-eGFP foci compared to *ssa1Δssa2Δ* strains, similar to wildtype (Fig. 4C). Therefore, the formation of TTR-eGFP foci is limited either by combined absence of both Ssa1 and Ssa2 or Ssa2 alone. Taken together, our data shows that in three separate techniques, deletion of both Ssa1 and Ssa2 increased the abundance of larger TTR-eGFP species compared to wildtype, suggesting that yeast Hsp70s help limit TTR-eGFP aggregation.

### Overexpression of Sis1 and Ssa1 have no significant change in TTR-eGFP sedimentation

We next asked how TTR-eGFP sedimentation would change upon chaperone overexpression. We focused on Sis1 and Ssa1 overexpression since Sse1 overexpression is toxic (Shaner, Trott et al. 2004). To evaluate the effects of Sis1 and Ssa1 overexpression on TTR-eGFP sedimentation, we utilized plasmids that increase Sis1 and Ssa1 stead levels by ∼5-fold and ∼1.5-fold, respectively, over wildtype (Buchholz 2025). We found that Sis1 overexpression in a wildtype strain did not alter TTR-eGFP sedimentation (Supplemental Fig. 10A). However, overexpression of Ssa1 in wildtype strains led to a modest, albeit not significant, increase in TTR-eGFP in Fraction 7 (Supplemental Fig. 10B). Given that the overexpression of Ssa1 and Sis1 has been linked with decreased heat shock factor 1 (Hsf1) activity (Krakowiak, Zheng et al. 2018, Feder, Ali et al. 2021), it is possible that the detection of TTR-eGFP in fraction 7 could be due to repression of the heat shock response, rather than a direct effect of Ssa1 on TTR-eGFP aggregation. However, the addition of excess Ssa1 or Sis1 in wildtype cells appears to have minimal effects on TTR-eGFP aggregation.

## Discussion

TTR is deposited into tissues as extracellular amyloid. Yet, several lines of evidence suggest that TTR is internalized and accumulates intracellularly (Misumi, Ando et al. 2013, Goncalves, Costelha et al. 2014, Lu, Mak et al. 2024). Here, we use a yeast model to explore how intracellular TTR is influenced by molecular chaperones. Our data show that yeast Hsp70s limit HMW species TTR-eGFP (Fig. 4), suggesting that Hsp70s play a role in mitigating further intracellular cellular aggregation. The similarity between intracellular yeast and human oligomers on native PAGE and the high conservation of Hsp70 function implies that similar Hsp70 mechanisms may occur in humans.

### Intracellular TTR forms HMW SDS-soluble aggregate species

In humans, TTR is a β-strand rich protein primarily present in the plasma as a tetramer. The rate-limiting step of TTR aggregation is the tetramer dissociation, as the subsequently released TTR monomers can rapidly misfold and aggregate (Colon and Kelly 1992, Lai, Colon et al. 1996, Johnson, Wiseman et al. 2005). TTR tetramer dissociation is slow and age-dependent, making studying TTR aggregation in mammalian models such as mice time-consuming (Goncalves, Costelha et al. 2014, Li, Lyu et al. 2018, Slamova, Adib et al. 2021). Using yeast, we not only detected TTR-eGFP aggregation within 24 hours of growth, but the aggregation profile is similar to the TTR observed in ATTR patient plasma (Pedretti, Wang et al. 2024). The similarity of the oligomer-like and HMW TTR-eGFP species seen in yeast (Fig. 1A) compared to the TTR in human patients suggests that TTR has an intrinsic propensity to form these HMW aggregate species.

We previously reported that TTR-eGFP exists in SDS-resistant forms in yeast (Knier, Davis et al. 2022). This study further clarifies that these SDS-resistant TTR species are dimer-like and tetramer-like species (Fig. 1A). The detectable TTR-eGFP aggregates are SDS-soluble as shown by native PAGE analysis (Fig. 1A), and our previous results by discontinuous sucrose gradients (Knier, Davis et al. 2022). These data indicate TTR-eGFP in yeast is not forming large SDS-resistant aggregates typically associated with β-strand rich and amyloid-forming proteins, such as yeast prions or other human aggregation-prone proteins (Kryndushkin, Alexandrov et al. 2003, Bagriantsev and Liebman 2004, Pigazzini, Lawrenz et al. 2021, Herod, Dyatel et al. 2022). Instead, TTR-eGFP may be forming more unstructured, amorphous-like aggregates such as those found in biomolecular condensates. Recent work has shown that proteins within biomolecular condensates can undergo liquid-to-solid transitions as a precursor to amyloid formation (Alberti and Hyman 2021, Zhang, Huang et al. 2023, Yan, Kuster et al. 2025). Given that GFP-tagged TTR can undergo liquid-liquid phase separation and subsequent liquid-to-solid transition into amyloid *in vitro* (Duan, Li et al. 2023), it is possible that TTR *in vivo* may form amorphous aggregates as an intermediate to amyloid.

### Hsp70s limits TTR-eGFP aggregation

Hsp70s role in limiting protein aggregation *in vivo* is well-documented in yeast, as various temperature sensitive model proteins and stress granule proteins form visible foci in *ssa1Δssa2Δ* strains under normal temperature conditions (Escusa-Toret, Vonk et al. 2013, Andersson, Eisele-Burger et al. 2021, Schneider, Ahmadpour et al. 2022, Rolli, Langridge et al. 2024, Buchholz, Martin et al. 2025). Here, we similarly find TTR-eGFP forms similar visible foci in *ssa1Δssa2Δ* strains. Additionally, native PAGE and sucrose gradients biochemically show that TTR-eGFP populations shift to HWM species (Fig. 4). The reduction of Sis1 and deletion of Sse1 resulted in a loss of TTR-eGFP in fraction 1 and a gain in fraction 7 in sucrose gradients (Fig. 2B and Fig. 3B), suggesting that a small portion of the TTR-eGFP population is shifting to HMW species in these strains. These data together suggest that yeast Hsp70s, Ssa1 and Ssa2, are important for limiting HMW SDS-soluble TTR-eGFP *in vivo,* while Sis1 and Sse1 many have more minor roles in limiting the formation.

Hsp70s are highly conserved and play critical roles in protein folding, disaggregation, and aging. In yeast, loss of Ssa1 and Ssa2 results in shortened replicative aging phenotypes (Oling, Eisele et al. 2014, Andersson, Eisele-Burger et al. 2021). Decline in Hsp70 levels with age is also observed in both mice and humans (Gutsmann-Conrad, Pahlavani et al. 1999, Njemini, Bautmans et al. 2011). However, the introduction of exogenous Hsp70 in mouse models improves lifespan (Bobkova, Evgen’ev et al. 2015), providing further support for the link between Hsp70 levels and aging. Given that ATTR is an age-related disease, it is possible that Hsp70s also play a role In TTR aggregation in humans. In ATTR patients, the Hsf1 transcription factor that activates Hsp70 expression in response to stress is increased, as well as Hsp70 itself (Santos, Magalhaes et al. 2008). In patient derived iPSC differentiated neuronal cells that internalize *ex vivo* TTR aggregates, the intracellular Hsp70, HspA1A, is also elevated (Leung, Nah et al. 2013). It is possible that the increase in Hsp70 may be a proteostasis response to the presence of intracellular TTR to limit further aggregation. Yet with aging, reduced Hps70 levels and compromised proteostasis may contribute to increased intracellular TTR aggregation and subsequent cellular decline.

### Possible mechanisms of yeast Hsp70 limiting TTR-eGFP aggregation

We suspect that the yeast Hsp70s and its co-chaperones could limit HMW SDS-soluble TTR-eGFP in a few ways. First, Hsp70 has been reported to have holdase activity, in which Hsp70 can bind to and sequester disease-associated proteins, such as tau and alpha synuclein, to prevent further aggregation (Patterson, Ward et al. 2011, Kundel, De et al. 2018, Tao, Berthet et al. 2021, Rutledge, Choy et al. 2022). The increase of TTR-eGFP size in *ssa1Δssa2Δ* could support holdase function. Additionally, the more subtle effects from the Hsp70 co-chaperones, Sis1 and Sse1, on TTR-eGFP suggests that Hsp70 may act independently of co-chaperones, potentially via holdase activity. Additionally, Sse1 holdase activity has also been well-documented (Oh, Chen et al. 1997, Goeckeler, Petruso et al. 2008, Polier, Dragovic et al. 2008, Polier, Hartl et al. 2010, Garcia, Nillegoda et al. 2017), and could act as an independent holdase as well, albeit with weaker effects relative to Hsp70s. Second, while the constitutive TTR-eGFP expression in our system makes measuring disaggregation difficult. However, if yeast Hsp70s play a role in TTR-eGFP disaggregation, Ssa2 may be the primary disaggregase because plasmid-expressed Ssa1 did not rescue TTR the percent of cells with foci in *ssa1Δssa2Δ* (Fig. 4C). Functional specialization between Ssa1 and Ssa2 has been previously observed, as Ssa1 and Ssa2 differ in curing of the yeast prion [*URE3*] (Schwimmer and Masison 2002) and in their efficiency of reactivating the model aggregating protein firefly luciferase *in vivo* (Hasin, Cusack et al. 2014). While Sse1 NEF activity often supports Hsp70 disaggregase activity, our data suggests that Sse1 NEF activity does not influence TTR-eGFP aggregation (Supplemental Fig. 8). Alternatively, the bulky size of Sse1 has been shown to structurally arrange Hsp70s on aggregates for disaggregation through entropic pulling (Wentink, Nillegoda et al. 2020), providing support that Sse1 could assist in Hsp70 disaggregation in a NEF-independent manner.

Third, the increased size of HMW SDS-soluble TTR-eGFP in *ssa1Δssa2Δ* may be a result of a dysregulated heat shock response and altered proteostasis. Yeast Hsp70s bind and sequester the Hsf1 in the cytoplasm (Krakowiak, Zheng et al. 2018), therefore repressing the heat shock response. Accordingly, the *ssa1Δssa2Δ* strain is in a constant state of elevated heat shock response state (Buchholz, Martin et al. 2025). Lastly, increased HMW SDS-soluble TTR-eGFP could result from chaperone-mediated sequestration of proteins in the *ssa1Δssa2Δ* background. The detection of abnormal protein aggregates in *ssa1Δssa2Δ* strains has been suggested not to reflect enhanced aggregation, but rather reflects the sequestration of aggregates by Btn2 and Hsp42 chaperones as a compensatory response to the loss of Hsp70 activity (Ho, Grousl et al. 2019). The sequestration of TTR-eGFP by Btn2 and Hsp42 into larger foci could also explain why the sucrose gradient assays were able to detect large changes in TTR-eGFP sedimentation.

The Hsp70 family has been shown to suppress the aggregation of several disease-associated proteins *in vitro* and *in vivo*, including those associated with cellular internalization such as Aβ (Evans, Wisen et al. 2006, Baughman, Clouser et al. 2018, Schneider, Gautam et al. 2021). Our findings provide *in vivo* evidence that Hsp70 limits TTR aggregation within the yeast cytosol and suggests Hsp70 functions as a central factor that mitigates intracellular TTR aggregation. The age-related decline in proteostasis and Hsp70 activity could enhance intracellular TTR aggregation in patients, leading to detrimental cellular effects. Despite our findings, the exact mechanism by which Hps70 limits TTR aggregation, either directly or indirectly, is still unknown. Defining how Hsp70 specifically limits TTR aggregation may be important for understanding how the chaperone influences intracellular aggregation in ATTR.

## Experimental Procedures

### Yeast Strains and Plasmids

All yeast strains (Supplementary Table 1) are in the 74-D694 background (Chernoff, Lindquist et al. 1995). Strains were cultured using standard synthetic media and growth procedures (Sherman, Fink et al. 1986) unless indicated. Introduction of plasmids are conducted using standard Lithium Acetate methods (Gietz and Woods 2002). Plasmids used in this study are listed in Supplementary Table 2. Human TTR cDNA used in this study lacks the 20 amino acid signal sequence and has a methionine to begin the TTR sequence (Derkatch, Uptain et al. 2004).

### Biochemical methods Native PAGE

Transformants were patched on solid selective media and grown overnight at 30°C. Fresh patches were used to inoculate 50 ml cultures in plasmid selective media, and cultures were grown at 30^°^C with shaking to log phase (OD₆₀₀ = 0.4–0.8). Lysates were isolated in 1X Tris-Glycine Native Sample Buffer (Invitrogen LC2673) containing 1% digitonin, supplemented with protease inhibitor cocktail and PMSF (Sigma-Aldrich P7626). Cultures were lysed using either glass bead lysis or a Cryolyser (Bertin Technologies) and precleared at 2,300 rpm for 1 minute. Approximately 20–35μg of total protein in 1X Tris-Glycine Native Sample Buffer was loaded onto either a Novex™ 4–20% Tris-Glycine Mini Protein Gel (1.0 mm, WedgeWell™ format; Invitrogen) or a NuPAGE™ 3–8% Tris-Acetate Mini Protein Gel (1.0–1.5 mm; Invitrogen). Separated proteins were transferred to PVDF and subjected to Western blot analysis. The iBright (Invitrogen) imaging system was used to visualize gels. ImageJ was used to calculate raw intensity values of all blots. For native PAGE gels quantification of TTR signal, the TTR intensity between 480 kDa and the top of the gel, or a region of interest at the top of the gel was divided by the total TTR intensity for the entire lane.

### Discontinuous Sucrose Gradient Sedimentation

Discontinuous sucrose gradient sedimentation was previously described (Dorweiler, Oddo et al. 2020, Knier, Davis et al. 2022). Briefly, lysates were isolated in 1X lysis buffer (50 mM Tris–HCl, 100 mM KCl, 20 mM MgCl₂•6H₂O, 2 mM EDTA, 10% glycerol; pH 7.5) supplemented with 1.22% yeast specific protease inhibitor cocktail (Sigma-Aldrich P8215) and 8.1µm PMSF (Sigma-Aldrich P7626) and precleared at 2,300 rpm for 1 minute. Approximately 2.2 mg of lysate was loaded onto a discontinuous (10%, 40%, 60%) sucrose gradient and spun at 16,000× *g* 4°C for 120 min in a swinging bucket rotor. Ten fractions of 180µL were collected, and the pellet was collected by adding 180µL of 1X Lysis Buffer and briefly vortexed. Samples were prepared by mixing equal volumes of fractions with 1X lysis buffer, followed by the addition of 2% SDS sample buffer. All samples were incubated at 95°C for 8 minutes prior to loading on a 10% SDS-PAGE. Membranes were subjected to Western blot analysis. For Triton X-100 studies, lysates were split into paired samples immediately after lysis. One sample was treated with 0.1% Triton X-100 (final concentration) and the other was left untreated. Treated and untreated lysates were incubated on ice for 10 minutes with occasional vortexing and inversion, then centrifuged at 2,300 rpm for 1 minute per usual for lysate clarification. To determine the percent TTR in each fraction, the TTR intensity of each 41 kDa band in each fraction was divided over the total TTR intensity from fractions 1–10 and the pellet.

### Spinning Disk Confocal Microscopy

Fresh patches were used to inoculate 4 mL of selective liquid media, and cultures were grown at 30°C with shaking to (OD₆₀₀ = 0.8–1.2). Field images were taken using confocal fluorescent microscopy, and a minimum of 200 cells analyzed per independent sample. Imaging was performed on a Zeiss AxioObserver microscope equipped with a CrestOptics Cicero spinning-disk confocal system (63X oil immersion objective, 1.44 NA) and an ORCA Flash CMOS camera. A z-series consisting of 12 optical sections was collected for each field of view. Images were acquired and processed using VisiView Software. All images shown as single z-stacks.

### Generation of Yeast Deletion Strains Single gene deletions

Single deletion strains were generated by replacing the open reading frames of target genes with the *HIS3* gene via homologous recombination according to Manogaran et al. (Manogaran, Hong et al. 2011). Briefly, primers (Supplementary Table 4) were designed to contain 40 bp flanking sequences to the target gene open reading frame and used to amplify the *HIS3* gene. The resulting product, containing *HIS3* flanked by target gene homology arms, was transformed into yeast using the lithium acetate method. Homologous recombination facilitated the replacement of the target gene’s open reading frame with the *HIS3* gene. Transformants were selected on synthetic complete medium lacking histidine. Correct integration was verified by diagnostic PCR using a forward primer located upstream of the 5′ homology region and a reverse primer within the *HIS3* gene. All primer sequences are listed in Supplementary Table 4.

### Generation of ssa1Δssa2Δ double deletion with Nat disruption

To generate the *ssa1Δssa2Δ* double deletion double deletion strain, primers designed with *SSA2* flank homology arms were used to amplify the nourseothricin resistance (*NAT1*) gene. PCR products were transformed into the *ssa1::HIS3* strain yeast strain using the lithium acetate method. Transformants were selected on YPD medium containing nourseothricin. Correct integration was confirmed by diagnostic PCR using primers located upstream of the 5′ homology region and within the NAT1 gene. Primer sequences are listed in Supplementary Table 4.

### Statistical analysis

Statistical analyses were performed using GraphPad Prism V. 10. All statistical tests are indicated in the figure legends. Statistical analyses for sucrose gradients were performed with a two-way repeated measures ANOVA, Geisser-Greenhouse correction, and a Sidak multiple comparisons test (* p < 0.05). Analyses were grouped as the following: Group #1: wildtype strains with and without Triton X-100, Group #2: *sis1*-*Δ* [*TETrSIS1*] with and without doxy, Group #3: wildtype strains, *sse2Δ, sse1Δ, ssa1Δ, ssa1Δ ssa2Δ,* wildtype with GPD-Sis1, and wildtype with GDP-Ssa1 (all comparisons made to wildtype control strain), and Group #4: *sse1Δ* with EV, *sse1Δ*, *sse1Δ* with Sse1^WT^, and *sse1Δ* with Sse1^G233D^ (All comparisons made to *sse1Δ* with EV control strain).

## Supporting information

This article contains supporting information.

## Acknowledgements.

The authors would like to thank Justin Hines (Lafayette College), Elizabeth Craig (UW-Madison) and Susan Liebman (UN-Reno) for strains, and Elizabeth Craig (UW-Madison) for antibodies used in these studies. Special thanks to Rosemary Stuart (Marquette) for technical expertise.

## Conflict of interest

The authors declare that they have no conflicts of interest with the contents of this article.

### Abbreviations

ATTR: transthyretin amyloidosis
AD: Alzheimer’s disease
TTR: transthyretin
Hsp70: heat shock protein 90
JDP: J-domain protein
Hsp110: heat shock protein 110
NEF: nucleotide exchange factor
HMW: high molecular weight
Hsf1: heat shock transcription factor 1
SDS: sodium dodecyl sulfate
Native PAGE: native polyacrylamide gel electrophoresis

## Literature Cited

1. Alberti, S. and A. A. Hyman (2021). “Biomolecular condensates at the nexus of cellular stress, protein aggregation disease and ageing.” Nat Rev Mol Cell Biol 22(3): 196–213.

2. Andersson, K., A. Olofsson, E. H. Nielsen, S. E. Svehag and E. Lundgren (2002). “Only amyloidogenic intermediates of transthyretin induce apoptosis.” Biochem Biophys Res Commun 294(2): 309–314.

3. Andersson, R., A. M. Eisele-Burger, S. Hanzen, K. Vielfort, D. Oling, F. Eisele, G. Johansson, T. Gustafsson, K. Kvint and T. Nystrom (2021). “Differential role of cytosolic Hsp70s in longevity assurance and protein quality control.” PLoS Genet 17(1): e1008951.

4. Andreasson, C., J. Fiaux, H. Rampelt, M. P. Mayer and B. Bukau (2008). “Hsp110 is a nucleotide-activated exchange factor for Hsp70.” J Biol Chem 283(14): 8877–8884.

5. Aron, R., T. Higurashi, C. Sahi and E. A. Craig (2007). “J-protein co-chaperone Sis1 required for generation of [RNQ+] seeds necessary for prion propagation.” Embo J 26(16): 3794–3803.

6. Bagriantsev, S. and S. W. Liebman (2004). “Specificity of prion assembly in vivo. [PSI+] and [PIN+] form separate structures in yeast.” J Biol Chem 279(49): 51042–51048.

7. Baughman, H. E. R., A. F. Clouser, R. E. Klevit and A. Nath (2018). “HspB1 and Hsc70 chaperones engage distinct tau species and have different inhibitory effects on amyloid formation.” J Biol Chem 293(8): 2687–2700.

8. Beton, J. G., J. Monistrol, A. Wentink, E. C. Johnston, A. J. Roberts, B. G. Bukau, B. W. Hoogenboom and H. R. Saibil (2022). “Cooperative amyloid fibre binding and disassembly by the Hsp70 disaggregase.” EMBO J 41(16): e110410.

9. Bobkova, N. V., M. Evgen’ev, D. G. Garbuz, A. M. Kulikov, A. Morozov, A. Samokhin, D. Velmeshev, N. Medvinskaya, I. Nesterova, A. Pollock and E. Nudler (2015). “Exogenous Hsp70 delays senescence and improves cognitive function in aging mice.” Proc Natl Acad Sci U S A 112(52): 16006–16011.

10. Buchholz, H. E., S. A. Martin, J. E. Dorweiler, C. M. Radtke, A. S. Knier, N. B. Beans and A. L. Manogaran (2025). “Hsp70 chaperones, Ssa1 and Ssa2, limit poly(A) binding protein aggregation.” Mol Biol Cell 36(6): ar66.

11. Buchholz, H. E., S. A. Martin, J. E. Dorweiler, C. M. Radtke, A. S. Knier, N. B. Beans and A. L. Manogaran (2025). “Hsp70 chaperones, Ssa1 and Ssa2, limit poly(A) binding protein aggregation.” Mol Biol Cell: mbcE25010027.

12. Buchholz, H. E., Martin, S.A., Dorweiler, J.E., Prosser, D.C., Sontag, E.M., Manogaran, A.L., (2025). “Stress granules and protein aggregates reveal intracellular resource competition.” bioRxiv.

13. Cheetham, M. E. and A. J. Caplan (1998). “Structure, function and evolution of DnaJ: conservation and adaptation of chaperone function.” Cell Stress Chaperones 3(1): 28–36.

14. Chernoff, Y. O., S. L. Lindquist, B. Ono, S. G. Inge-Vechtomov and S. W. Liebman (1995). “Role of the chaperone protein Hsp104 in propagation of the yeast prion-like factor [psi+].” Science 268(5212): 880–884.

15. Colon, W. and J. W. Kelly (1992). “Partial denaturation of transthyretin is sufficient for amyloid fibril formation in vitro.” Biochemistry 31(36): 8654–8660.

16. Dasari, A. K. R., R. M. Hughes, S. Wi, I. Hung, Z. Gan, J. W. Kelly and K. H. Lim (2019). “Transthyretin Aggregation Pathway toward the Formation of Distinct Cytotoxic Oligomers.” Sci Rep 9(1): 33.

17. Dasari, A. K. R., I. Hung, Z. Gan and K. H. Lim (2019). “Two distinct aggregation pathways in transthyretin misfolding and amyloid formation.” Biochim Biophys Acta Proteins Proteom 1867(3): 344–349.

18. De Kimpe, L., E. S. van Haastert, A. Kaminari, R. Zwart, H. Rutjes, J. J. Hoozemans and W. Scheper (2013). “Intracellular accumulation of aggregated pyroglutamate amyloid beta: convergence of aging and Abeta pathology at the lysosome.” Age (Dordr) 35(3): 673–687.

19. Derkatch, I. L., S. M. Uptain, T. F. Outeiro, R. Krishnan, S. L. Lindquist and S. W. Liebman (2004). “Effects of Q/N-rich, polyQ, and non-polyQ amyloids on the de novo formation of the [PSI+] prion in yeast and aggregation of Sup35 in vitro.” Proc Natl Acad Sci U S A 101(35): 12934–12939.

20. Dorweiler, J. E., M. J. Oddo, D. R. Lyke, J. A. Reilly, B. T. Wisniewski, E. E. Davis, A. M. Kuborn, S. J. Merrill and A. L. Manogaran (2020). “The actin cytoskeletal network plays a role in yeast prion transmission and contributes to prion stability.” Mol Microbiol.

21. Duan, G., Y. Li, M. Ye, H. Liu, N. Wang and S. Luo (2023). “The Regulatory Mechanism of Transthyretin Irreversible Aggregation through Liquid-to-Solid Phase Transition.” Int J Mol Sci 24(4).

22. Escusa-Toret, S., W. I. Vonk and J. Frydman (2013). “Spatial sequestration of misfolded proteins by a dynamic chaperone pathway enhances cellular fitness during stress.” Nat Cell Biol 15(10): 1231–1243.

23. Evans, C. G., S. Wisen and J. E. Gestwicki (2006). “Heat shock proteins 70 and 90 inhibity early stages of amyloid beta-(1-42) aggregation in vitro.” J Biol Chem 281(44): 33182–33191.

24. Fan, C. Y., S. Lee and D. M. Cyr (2003). “Mechanisms for regulation of Hsp70 function by Hsp40.” Cell Stress Chaperones 8(4): 309–316.

25. Feder, Z. A., A. Ali, A. Singh, J. Krakowiak, X. Zheng, V. P. Bindokas, D. Wolfgeher, S. J. Kron and D. Pincus (2021). “Subcellular localization of the J-protein Sis1 regulates the heat shock response.” J Cell Biol 220(1).

26. Gallego Villarejo, L., L. Bachmann, D. Marks, M. Brachthauser, A. Geidies and T. Muller (2022). “Role of Intracellular Amyloid beta as Pathway Modulator, Biomarker, and Therapy Target.” Int J Mol Sci 23(9).

27. Gao, L., X. Xie, P. Liu and J. Jin (2022). “High-avidity binding drives nucleation of amyloidogenic transthyretin monomer.” JCI Insight 7(7).

28. Gao, X., M. Carroni, C. Nussbaum-Krammer, A. Mogk, N. B. Nillegoda, A. Szlachcic, D. L. Guilbride, H. R. Saibil, M. P. Mayer and B. Bukau (2015). “Human Hsp70 Disaggregase Reverses Parkinson’s-Linked alpha-Synuclein Amyloid Fibrils.” Mol Cell 59(5): 781–793.

29. Garcia, V. M., N. B. Nillegoda, B. Bukau and K. A. Morano (2017). “Substrate binding by the yeast Hsp110 nucleotide exchange factor and molecular chaperone Sse1 is not obligate for its biological activities.” Mol Biol Cell 28(15): 2066–2075.

30. Gietz, R. D. and R. A. Woods (2002). “Transformation of yeast by lithium acetate/single-stranded carrier DNA/polyethylene glycol method.” Methods Enzymol 350: 87–96.

31. Goeckeler, J. L., A. P. Petruso, J. Aguirre, C. C. Clement, G. Chiosis and J. L. Brodsky (2008). “The yeast Hsp110, Sse1p, exhibits high-affinity peptide binding.” FEBS Lett 582(16): 2393–2396.

32. Goncalves, N. P., S. Costelha and M. J. Saraiva (2014). “Glial cells in familial amyloidotic polyneuropathy.” Acta Neuropathol Commun 2: 177.

33. Gutsmann-Conrad, A., M. A. Pahlavani, A. R. Heydari and A. Richardson (1999). “Expression of heat shock protein 70 decreases with age in hepatocytes and splenocytes from female rats.” Mech Ageing Dev 107(3): 255–270.

34. Hasin, N., S. A. Cusack, S. S. Ali, D. A. Fitzpatrick and G. W. Jones (2014). “Global transcript and phenotypic analysis of yeast cells expressing Ssa1, Ssa2, Ssa3 or Ssa4 as sole source of cytosolic Hsp70-Ssa chaperone activity.” BMC Genomics 15(1): 194.

35. Henze, A., T. Homann, M. Serteser, O. Can, O. Sezgin, A. Coskun, I. Unsal, F. J. Schweigert and A. Ozpinar (2015). “Post-translational modifications of transthyretin affect the triiodonine-binding potential.” J Cell Mol Med 19(2): 359–370.

36. Herod, S. G., A. Dyatel, S. Hodapp, M. Jovanovic and L. E. Berchowitz (2022). “Clearance of an amyloid-like translational repressor is governed by 14-3-3 proteins.” Cell Rep 39(5): 110753.

37. Hivare, P., K. Mujmer, G. Swarup, S. Gupta and D. Bhatia (2023). “Endocytic pathways of pathogenic protein aggregates in neurodegenerative diseases.” Traffic 24(10): 434–452.

38. Ho, B., A. Baryshnikova and G. W. Brown (2018). “Unification of Protein Abundance Datasets Yields a Quantitative Saccharomyces cerevisiae Proteome.” Cell Syst 6(2): 192–205 e193.

39. Ho, C. T., T. Grousl, O. Shatz, A. Jawed, C. Ruger-Herreros, M. Semmelink, R. Zahn, K. Richter, B. Bukau and A. Mogk (2019). “Cellular sequestrases maintain basal Hsp70 capacity ensuring balanced proteostasis.” Nat Commun 10(1): 4851.

40. Hoshino, T., N. Murao, T. Namba, M. Takehara, H. Adachi, M. Katsuno, G. Sobue, T. Matsushima, T. Suzuki and T. Mizushima (2011). “Suppression of Alzheimer’s disease-related phenotypes by expression of heat shock protein 70 in mice.” J Neurosci 31(14): 5225–5234.

41. Jayaweera, S. W., M. Sahin, F. Lundkvist, A. Leven, L. Tereenstra, J. Backman, A. Bachhar, F. Bano, I. Anan and A. Olofsson (2025). “Misfolding of transthyretin in vivo is controlled by the redox environment and macromolecular crowding.” J Biol Chem 301(1): 108031.

42. Johnson, S. M., R. L. Wiseman, Y. Sekijima, N. S. Green, S. L. Adamski-Werner and J. W. Kelly (2005). “Native state kinetic stabilization as a strategy to ameliorate protein misfolding diseases: a focus on the transthyretin amyloidoses.” Acc Chem Res 38(12): 911–921.

43. Kampinga, H. H. and E. A. Craig (2010). “The HSP70 chaperone machinery: J proteins as drivers of functional specificity.” Nat Rev Mol Cell Biol 11(8): 579–592.

44. Kingsbury, J. S., T. M. Laue, E. S. Klimtchuk, R. Theberge, C. E. Costello and L. H. Connors (2008). “The modulation of transthyretin tetramer stability by cysteine 10 adducts and the drug diflunisal. Direct analysis by fluorescence-detected analytical ultracentrifugation.” J Biol Chem 283(18): 11887–11896.

45. Knier, A. S., E. E. Davis, H. E. Buchholz, J. E. Dorweiler, L. E. Flannagan and A. L. Manogaran (2022). “The yeast molecular chaperone, Hsp104, influences Transthyretin (TTR) aggregate formation.” Front. Mol. Neurosci.

46. Krakowiak, J., X. Zheng, N. Patel, Z. A. Feder, J. Anandhakumar, K. Valerius, D. S. Gross, A. S. Khalil and D. Pincus (2018). “Hsf1 and Hsp70 constitute a two-component feedback loop that regulates the yeast heat shock response.” Elife 7.

47. Kryndushkin, D. S., I. M. Alexandrov, M. D. Ter-Avanesyan and V. V. Kushnirov (2003). “Yeast [PSI+] prion aggregates are formed by small Sup35 polymers fragmented by Hsp104.” J Biol Chem 278(49): 49636–49643.

48. Kundel, F., S. De, P. Flagmeier, M. H. Horrocks, M. Kjaergaard, S. L. Shammas, S. E. Jackson, C. M. Dobson and D. Klenerman (2018). “Hsp70 Inhibits the Nucleation and Elongation of Tau and Sequesters Tau Aggregates with High Affinity.” ACS Chem Biol 13(3): 636–646.

49. LaFerla, F. M., K. N. Green and S. Oddo (2007). “Intracellular amyloid-beta in Alzheimer’s disease.” Nat Rev Neurosci 8(7): 499–509.

50. Lai, Z., W. Colon and J. W. Kelly (1996). “The acid-mediated denaturation pathway of transthyretin yields a conformational intermediate that can self-assemble into amyloid.” Biochemistry 35(20): 6470–6482.

51. Lee do, H., M. Y. Sherman and A. L. Goldberg (2016). “The requirements of yeast Hsp70 of SSA family for the ubiquitin-dependent degradation of short-lived and abnormal proteins.” Biochem Biophys Res Commun 475(1): 100–106.

52. Leung, A., S. K. Nah, W. Reid, A. Ebata, C. M. Koch, S. Monti, J. C. Genereux, R. L. Wiseman, B. Wolozin, L. H. Connors, J. L. Berk, D. C. Seldin, G. Mostoslavsky, D. N. Kotton and G. J. Murphy (2013). “Induced pluripotent stem cell modeling of multisystemic, hereditary transthyretin amyloidosis.” Stem Cell Reports 1(5): 451–463.

53. Li, X., Y. Lyu, J. Shen, Y. Mu, L. Qiang, L. Liu, K. Araki, B. P. Imbimbo, K. I. Yamamura, S. Jin and Z. Li (2018). “Amyloid deposition in a mouse model humanized at the transthyretin and retinol-binding protein 4 loci.” Lab Invest 98(4): 512–524.

54. Lim, A., T. Prokaeva, M. E. McComb, P. B. O’Connor, R. Theberge, L. H. Connors, M. Skinner and C. E. Costello (2002). “Characterization of transthyretin variants in familial transthyretin amyloidosis by mass spectrometric peptide mapping and DNA sequence analysis.” Anal Chem 74(4): 741–751.

55. Liu, R. Q., Q. H. Zhou, S. R. Ji, Q. Zhou, D. Feng, Y. Wu and S. F. Sui (2010). “Membrane localization of beta-amyloid 1-42 in lysosomes: a possible mechanism for lysosome labilization.” J Biol Chem 285(26): 19986–19996.

56. Lu, J. Q., G. Mak, S. Grant and S. K. Baker (2024). “Hereditary Transthyretin Amyloidosis Neuropathy with Intracellular Amyloidosis and Inclusions.” Can J Neurol Sci: 1–3.

57. Magrane, J., R. C. Smith, K. Walsh and H. W. Querfurth (2004). “Heat shock protein 70 participates in the neuroprotective response to intracellularly expressed beta-amyloid in neurons.” J Neurosci 24(7): 1700–1706.

58. Manogaran, A. L., J. Y. Hong, J. Hufana, J. Tyedmers, S. Lindquist and S. W. Liebman (2011). “Prion formation and polyglutamine aggregation are controlled by two classes of genes.” PLoS Genet 7(5): e1001386.

59. Matsumoto, R., K. Akama, R. Rakwal and H. Iwahashi (2005). “The stress response against denatured proteins in the deletion of cytosolic chaperones SSA1/2 is different from heat-shock response in Saccharomyces cerevisiae.” BMC Genomics 6: 141.

60. Misumi, Y., Y. Ando, N. P. Goncalves and M. J. Saraiva (2013). “Fibroblasts endocytose and degrade transthyretin aggregates in transthyretin-related amyloidosis.” Lab Invest 93(8): 911–920.

61. Murakami, T., Y. Ito, K. Sango, K. Watabe and Y. Sunada (2023). “Human transthyretin gene expression is markedly increased in repair Schwann cells in an in vitro model of hereditary transthyretin amyloidosis.” Neurochem Int 164: 105507.

62. Nachman, E., A. S. Wentink, K. Madiona, L. Bousset, T. Katsinelos, K. Allinson, H. Kampinga, W. A. McEwan, T. R. Jahn, R. Melki, A. Mogk, B. Bukau and C. Nussbaum-Krammer (2020). “Disassembly of Tau fibrils by the human Hsp70 disaggregation machinery generates small seeding-competent species.” J Biol Chem 295(28): 9676–9690.

63. Njemini, R., I. Bautmans, O. O. Onyema, K. Van Puyvelde, C. Demanet and T. Mets (2011). “Circulating heat shock protein 70 in health, aging and disease.” BMC Immunol 12: 24.

64. Nollen, E. A. and R. I. Morimoto (2002). “Chaperoning signaling pathways: molecular chaperones as stress-sensing ’heat shock’ proteins.” J Cell Sci 115(Pt 14): 2809–2816.

65. Oh, H. J., X. Chen and J. R. Subjeck (1997). “Hsp110 protects heat-denatured proteins and confers cellular thermoresistance.” J Biol Chem 272(50): 31636–31640.

66. Oling, D., F. Eisele, K. Kvint and T. Nystrom (2014). “Opposing roles of Ubp3-dependent deubiquitination regulate replicative life span and heat resistance.” EMBO J 33(7): 747–761.

67. Patterson, K. R., S. M. Ward, B. Combs, K. Voss, N. M. Kanaan, G. Morfini, S. T. Brady, T. C. Gamblin and L. I. Binder (2011). “Heat shock protein 70 prevents both tau aggregation and the inhibitory effects of preexisting tau aggregates on fast axonal transport.” Biochemistry 50(47): 10300–10310.

68. Pedretti, R., L. Wang, A. Yakubovska, Q. S. Zhang, B. Nguyen, J. L. Grodin, A. Masri and L. Saelices (2024). “Structure-Based Probe Reveals the Presence of Large Transthyretin Aggregates in Plasma of ATTR Amyloidosis Patients.” JACC Basic Transl Sci 9(9): 1088–1100.

69. Pigazzini, M. L., M. Lawrenz, A. Margineanu, G. S. Kaminski Schierle and J. Kirstein (2021). “An Expanded Polyproline Domain Maintains Mutant Huntingtin Soluble in vivo and During Aging.” Front Mol Neurosci 14: 721749.

70. Polier, S., Z. Dragovic, F. U. Hartl and A. Bracher (2008). “Structural basis for the cooperation of Hsp70 and Hsp110 chaperones in protein folding.” Cell 133(6): 1068–1079.

71. Polier, S., F. U. Hartl and A. Bracher (2010). “Interaction of the Hsp110 molecular chaperones from S. cerevisiae with substrate protein.” J Mol Biol 401(5): 696–707.

72. Raviol, H., H. Sadlish, F. Rodriguez, M. P. Mayer and B. Bukau (2006). “Chaperone network in the yeast cytosol: Hsp110 is revealed as an Hsp70 nucleotide exchange factor.” EMBO J 25(11): 2510–2518.

73. Reixach, N., S. Deechongkit, X. Jiang, J. W. Kelly and J. N. Buxbaum (2004). “Tissue damage in the amyloidoses: Transthyretin monomers and nonnative oligomers are the major cytotoxic species in tissue culture.” Proc Natl Acad Sci U S A 101(9): 2817–2822.

74. Rolli, S., C. A. Langridge and E. M. Sontag (2024). “Clearing the JUNQ: the molecular machinery for sequestration, localization, and degradation of the JUNQ compartment.” Front Mol Biosci 11: 1427542.

75. Rutledge, B. S., W. Y. Choy and M. L. Duennwald (2022). "Folding or holding?-Hsp70 and Hsp90 chaperoning of misfolded proteins in neurodegenerative disease.” J Biol Chem 298(5): 101905.

76. Santos, S. D., R. Fernandes and M. J. Saraiva (2010). “The heat shock response modulates transthyretin deposition in the peripheral and autonomic nervous systems.” Neurobiol Aging 31(2): 280–289.

77. Santos, S. D., J. Magalhaes and M. J. Saraiva (2008). “Activation of the heat shock response in familial amyloidotic polyneuropathy.” J Neuropathol Exp Neurol 67(5): 449–455.

78. Schneider, K. L., D. Ahmadpour, K. S. Keuenhof, A. M. Eisele-Burger, L. L. Berglund, F. Eisele, R. Babazadeh, J. L. Hoog, T. Nystrom and P. O. Widlund (2022). “Using reporters of different misfolded proteins reveals differential strategies in processing protein aggregates.” J Biol Chem 298(11): 102476.

79. Schneider, M. M., S. Gautam, T. W. Herling, E. Andrzejewska, G. Krainer, A. M. Miller, V. A. Trinkaus, Q. A. E. Peter, F. S. Ruggeri, M. Vendruscolo, A. Bracher, C. M. Dobson, F. U. Hartl and T. P. J. Knowles (2021). “The Hsc70 disaggregation machinery removes monomer units directly from alpha-synuclein fibril ends.” Nat Commun 12(1): 5999.

80. Schwimmer, C. and D. C. Masison (2002). “Antagonistic interactions between yeast [PSI(+)] and [URE3] prions and curing of [URE3] by Hsp70 protein chaperone Ssa1p but not by Ssa2p.” Mol Cell Biol 22(11): 3590–3598.

81. Shaner, L., A. Trott, J. L. Goeckeler, J. L. Brodsky and K. A. Morano (2004). “The function of the yeast molecular chaperone Sse1 is mechanistically distinct from the closely related hsp70 family.” J Biol Chem 279(21): 21992–22001.

82. Shaner, L., H. Wegele, J. Buchner and K. A. Morano (2005). “The yeast Hsp110 Sse1 functionally interacts with the Hsp70 chaperones Ssa and Ssb.” J Biol Chem 280(50): 41262–41269.

83. Sherman, F., G. R. Fink and J. B. Hicks (1986). *Methods in Yeast Genetics*. Plainview, New York, Cold Spring Harbor Lab.

84. Shorter, J. (2011). “The mammalian disaggregase machinery: Hsp110 synergizes with Hsp70 and Hsp40 to catalyze protein disaggregation and reactivation in a cell-free system.” PLoS One 6(10): e26319.

85. Slamova, I., R. Adib, S. Ellmerich, M. R. Golos, J. A. Gilbertson, N. Botcher, D. Canetti, G. W. Taylor, N. Rendell, G. A. Tennent, G. Verona, R. Porcari, P. P. Mangione, J. D. Gillmore, M. B. Pepys, V. Bellotti, P. N. Hawkins, R. Al-Shawi and J. P. Simons (2021). “Plasmin activity promotes amyloid deposition in a transgenic model of human transthyretin amyloidosis.” Nat Commun 12(1): 7112.

86. Sorgjerd, K., T. Klingstedt, M. Lindgren, K. Kagedal and P. Hammarstrom (2008). “Prefibrillar transthyretin oligomers and cold stored native tetrameric transthyretin are cytotoxic in cell culture.” Biochem Biophys Res Commun 377(4): 1072–1078.

87. Sousa, M. M., I. Cardoso, R. Fernandes, A. Guimaraes and M. J. Saraiva (2001). “Deposition of transthyretin in early stages of familial amyloidotic polyneuropathy: evidence for toxicity of nonfibrillar aggregates.” Am J Pathol 159(6): 1993–2000.

88. Sousa, M. M. and M. J. Saraiva (2001). “Internalization of transthyretin. Evidence of a novel yet unidentified receptor-associated protein (RAP)-sensitive receptor.” J Biol Chem 276(17): 14420–14425.

89. Tao, J., A. Berthet, Y. R. Citron, P. L. Tsiolaki, R. Stanley, J. E. Gestwicki, D. A. Agard and L. McConlogue (2021). “Hsp70 chaperone blocks alpha-synuclein oligomer formation via a novel engagement mechanism.” J Biol Chem 296: 100613.

90. Verma, M., A. Girdhar, B. Patel, N. K. Ganguly, R. Kukreti and V. Taneja (2018). “Q-Rich Yeast Prion [PSI(+)] Accelerates Aggregation of Transthyretin, a Non-Q-Rich Human Protein.” Front Mol Neurosci 11: 75.

91. Wentink, A. S., N. B. Nillegoda, J. Feufel, G. Ubartaite, C. P. Schneider, P. De Los Rios, J. Hennig, A. Barducci and B. Bukau (2020). “Molecular dissection of amyloid disaggregation by human HSP70.” Nature 587(7834): 483–488.

92. Werner-Washburne, M., D. E. Stone and E. A. Craig (1987). “Complex interactions among members of an essential subfamily of hsp70 genes in Saccharomyces cerevisiae.” Mol Cell Biol 7(7): 2568–2577.

93. Wyszkowski, H., A. Janta, W. Sztangierska, I. Obuchowski, T. Chamera, A. Klosowska and K. Liberek (2021). “Class-specific interactions between Sis1 J-domain protein and Hsp70 chaperone potentiate disaggregation of misfolded proteins.” Proc Natl Acad Sci U S A 118(49).

94. Yan, X., D. Kuster, P. Mohanty, J. Nijssen, K. Pombo-Garcia, J. Garcia Morato, A. Rizuan, T. M. Franzmann, A. Sergeeva, A. M. Ly, F. Liu, P. M. Passos, L. George, S. H. Wang, J. Shenoy, H. L. Danielson, B. Ozguney, A. Honigmann, Y. M. Ayala, N. L. Fawzi, D. W. Dickson, W. Rossoll, J. Mittal, S. Alberti and A. A. Hyman (2025). “Intra-condensate demixing of TDP-43 inside stress granules generates pathological aggregates.” Cell.

95. Zhang, Z., G. Huang, Z. Song, A. J. Gatch and F. Ding (2023). “Amyloid Aggregation and Liquid-Liquid Phase Separation from the Perspective of Phase Transitions.” J Phys Chem B 127(28): 6241–6250.

