## Supporting Information for "Intracellular aggregation of the transthyretin protein is limited by Hsp70 chaperones"

**Running title:** Hsp70 limits TTR aggregation

**Keywords**: protein aggregation, molecular chaperones, Hsp70, disaggregation, proteostasis, transthyretin

### Supporting Experimental Procedures

#### *Serial Dilution Plating Assay*

Transformants were patched on solid selective media and grown overnight at 30°C. Fresh patches were used to inoculate 4 ml cultures in plasmid selective media. Strains were grown at 30˚C, normalized for cell density, spotted onto selective media in 5-fold serial dilutions, and incubated at 30˚C for several days.

#### *Generation of Sse1$^{WT}$ and Sse1$^{G233D}$ plasmid*

The *SSE1* gene, including its endogenous promoter, coding sequence, and terminator, was PCR-amplified from genomic DNA from the 74D-694 background using gene-specific primers listed in Supplementary Table 3. The purified PCR product was introduced into the pGEM-T Easy vector (Promega) and verified by *EcoRI* and *BglII* digestion. Site-directed mutagenesis using the Q5® Site-Directed Mutagenesis Kit (New England Biolabs, E0554) was performed to generate the Sse1$^{G233D}$ point mutation. Clones were sequenced for verification. Wildtype and mutant *SSE1* constructs were introduced into the pRS314 yeast expression plasmid by *ClaI* digestion. Successful clones were confirmed by colony PCR and/or restriction digest analysis.

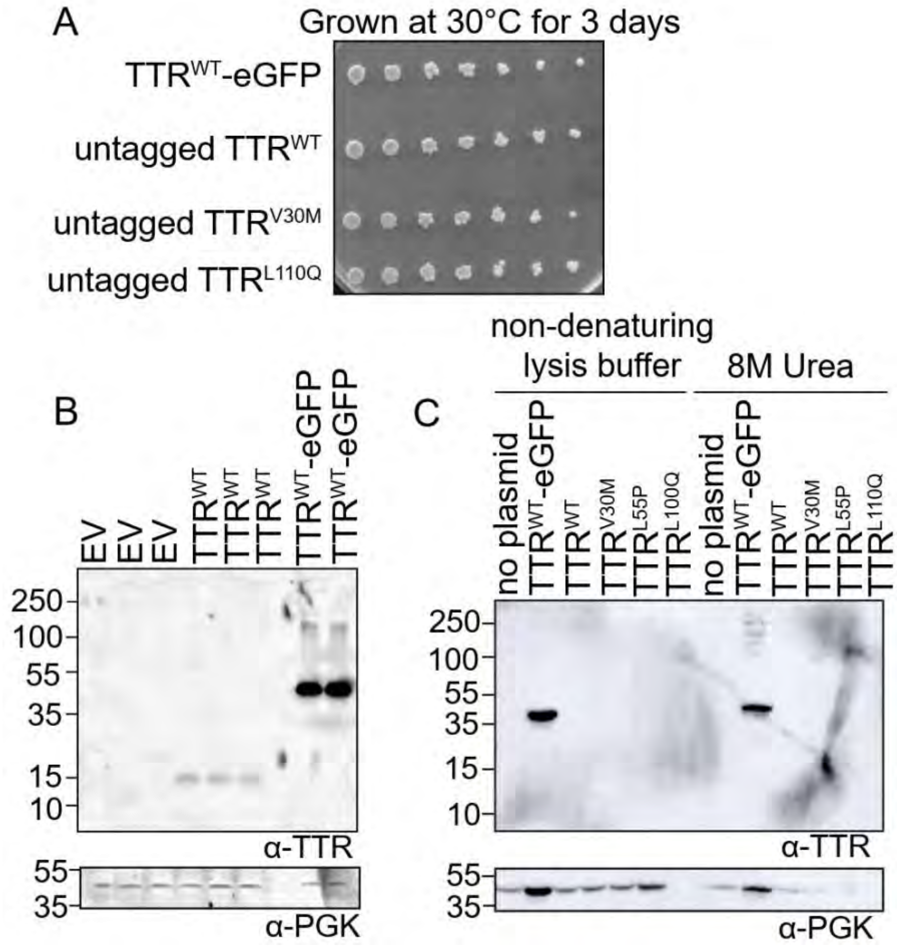


**Supplemental Figure 1. Untagged TTR exhibits low to no detection compared to TTR-eGFP.** A) 74-D694 wildtype strains (D230) were transformed with TTR$^{WT}$-eGFP (p3146), untagged TTR$^{WT}$ (p3235), untagged TTR$^{V30M}$ (p3240), and untagged TTR$^{L110Q}$ (p3244). Cells were serially diluted 5-fold, spotted on SD-Ura plates, and grown at 30°C for three days. Shown is a representative image from two independent trials. B) Wildtype strains were transformed with empty vector (EV, p3299), untagged TTR$^{WT}$ (p3301), and TTR$^{WT}$-eGFP. Samples were analyzed by SDS-PAGE and immunoblotted with polyclonal anti-TTR or anti-PGK antibodies. C) Wildtype strains were transformed with TTR-eGFP, untagged TTR$^{WT}$, untagged TTR$^{V30M}$, untagged TTR$^{L55P}$ (p3242), and untagged TTR$^{L110Q}$. Cultures were split in two and lysates isolated either in standard non-denaturing 1X lysis buffer or in the presence of 50mM HEPES pH 7.4 and 8M urea. Samples were analyzed by SDS-PAGE and immunoblotted with the indicated monoclonal anti-TTR or anti-PGK antibodies.

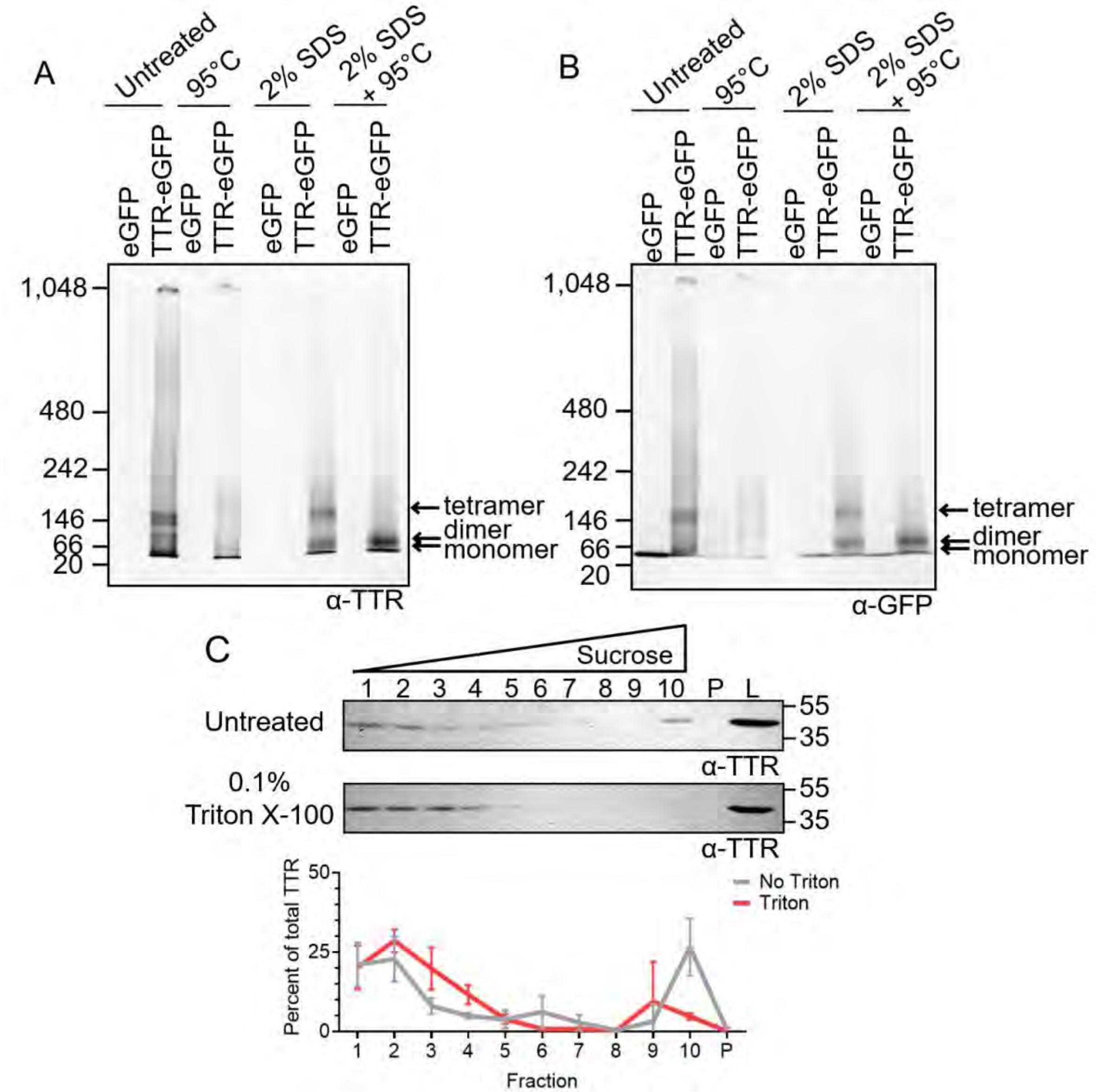


**Supplemental Figure 2. Tris-acetate native PAGE analysis and triton X-100 solubility of TTR-eGFP.** Wildtype strains were transformed eGFP or TTR-eGFP. Lysates were untreated, incubated at 95°C for 8 minutes, treated with 2% SDS, or 2% SDS and incubated at 95°C for 8 minutes. Lysates were analyzed by 3-8% tris-acetate native PAGE and immunoblotted with A) polyclonal anti-TTR or B) anti-GFP. C) Lysates from wildtype strains transformed TTR-eGFP were split and either left untreated or treated with 0.1% Triton X-100 for 10 minutes prior to loading on the sucrose gradient. All fractions were analyzed by SDS-PAGE and immunoblotted with a monoclonal anti-TTR antibody. Shown is a representative of three independent biological trials. The TTR signal from each fraction was normalized to the combined TTR signal in fractions 1-10 and pellet and graphed as the mean ± SD. Statistical analyses were performed with a two-way repeated measures ANOVA, Geisser-Greenhouse correction, and a Sidak multiple comparisons test (* $p < 0.05$). No comparisons are significant.

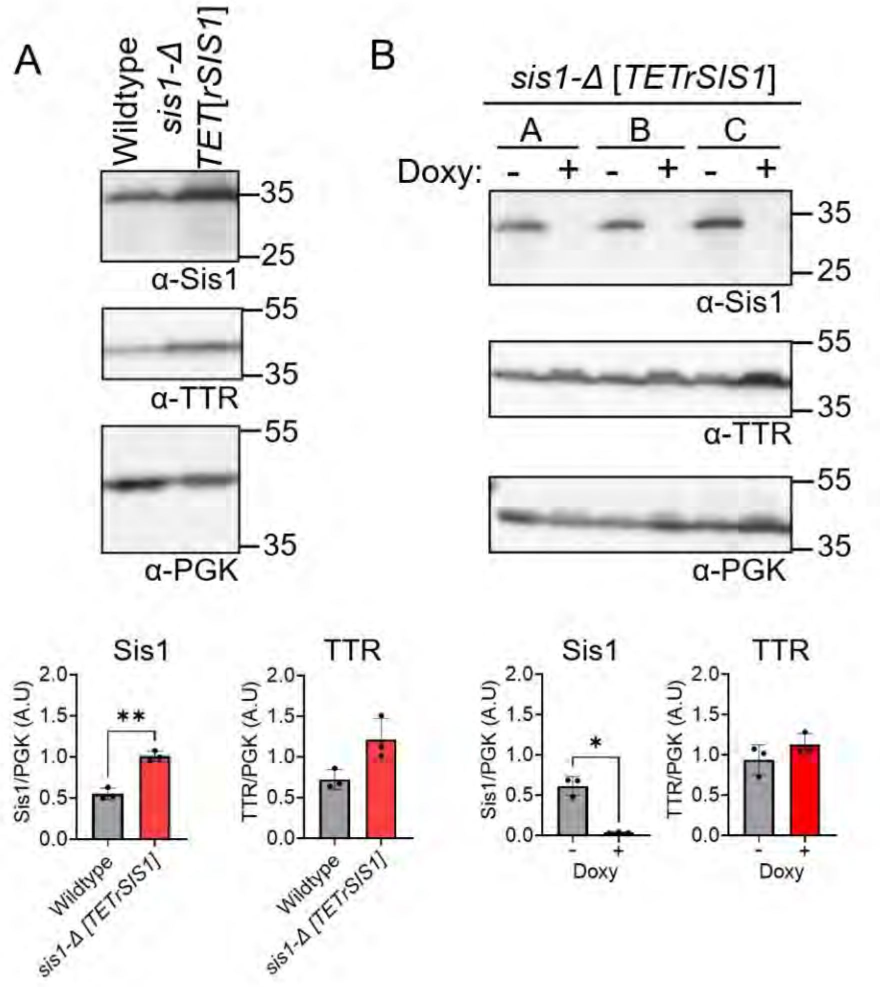


**Supplemental Figure 3. Doxycycline reduces tetracycline-repressible Sis1 levels by approximately 17-fold.** A) Wildtype and *sis1-Δ* [*TETrSIS1*] strains were transformed with TTR-eGFP. Untreated lysates were analyzed by SDS-PAGE and immunoblotted with the indicated antibodies. Statistical analyses were performed with an unpaired t-test (** $p < 0.01$). B) *sis1-Δ* [*TETrSIS1*] strains were transformed with TTR-eGFP. Cultures were split into untreated and treated with 10 µg/mL doxycycline overnight. Lysates from three independent replicates (A-C) were analyzed by SDS-PAGE followed by immunoblotted with the indicated antibodies. Statistical analyses were performed using paired t-test (* $p < 0.05$). All graphs have dots as independent trials and graphed as means ± SD. Unlabeled comparisons are not significant.

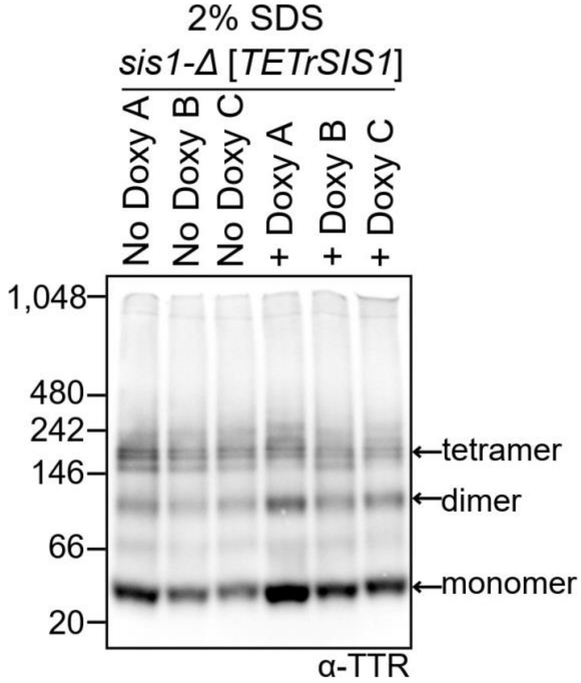


**Supplemental Figure 4. SDS treated *sis1-Δ* [*TETrSIS1*] shows no difference in TTR-eGFP on native PAGE.** *sis1-Δ* [*TETrSIS1*] cultures expressing TTR-eGFP were split and either left untreated or treated with 10 µg/mL doxycycline overnight. Lysates were treated with 2% SDS prior to 4–20% tris-glycine native PAGE and immunoblotted in polyclonal anti-TTR antibody. Estimated monomeric, dimeric, and tetrameric TTR-eGFP sizes are labeled.

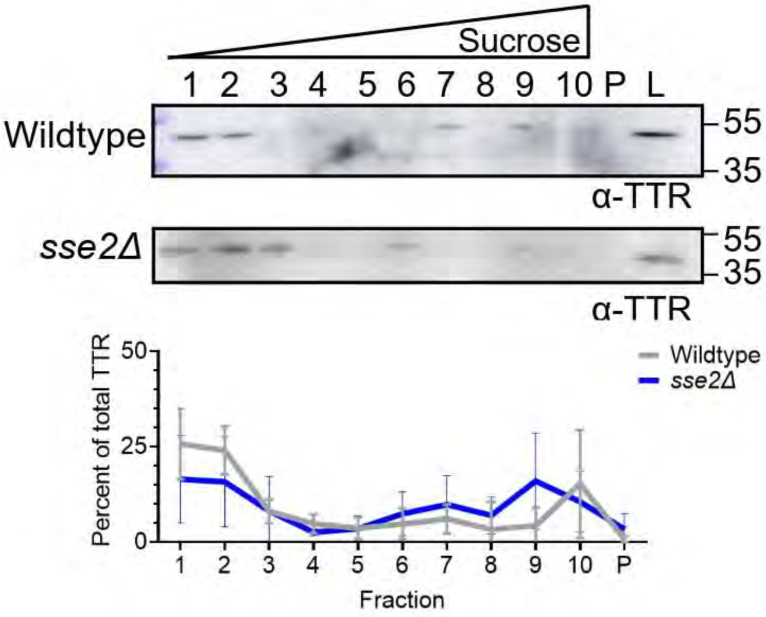


**Supplemental Figure 5. Deletion of Sse2 does not alter TTR-eGFP sedimentation.** Lysates from *sse2Δ* (M650) cultures expressing TTR-eGFP were loaded onto discontinuous sucrose gradients (10%, 40%, and 60%). Individual fractions, pelleted protein (P), and whole-cell lysates (L) were analyzed by SDS-PAGE and immunoblotted with a monoclonal anti-TTR antibody. Shown is a representative blot from four independent trials. The TTR signal from each fraction was normalized to the combined TTR signal in fractions 1-10 and pellet and graphed as mean ± SD. Statistical analyses were performed with a two-way repeated measures ANOVA, Geisser-Greenhouse correction, and a Sidak multiple comparisons test (* $p < 0.05$). No comparisons are significant.

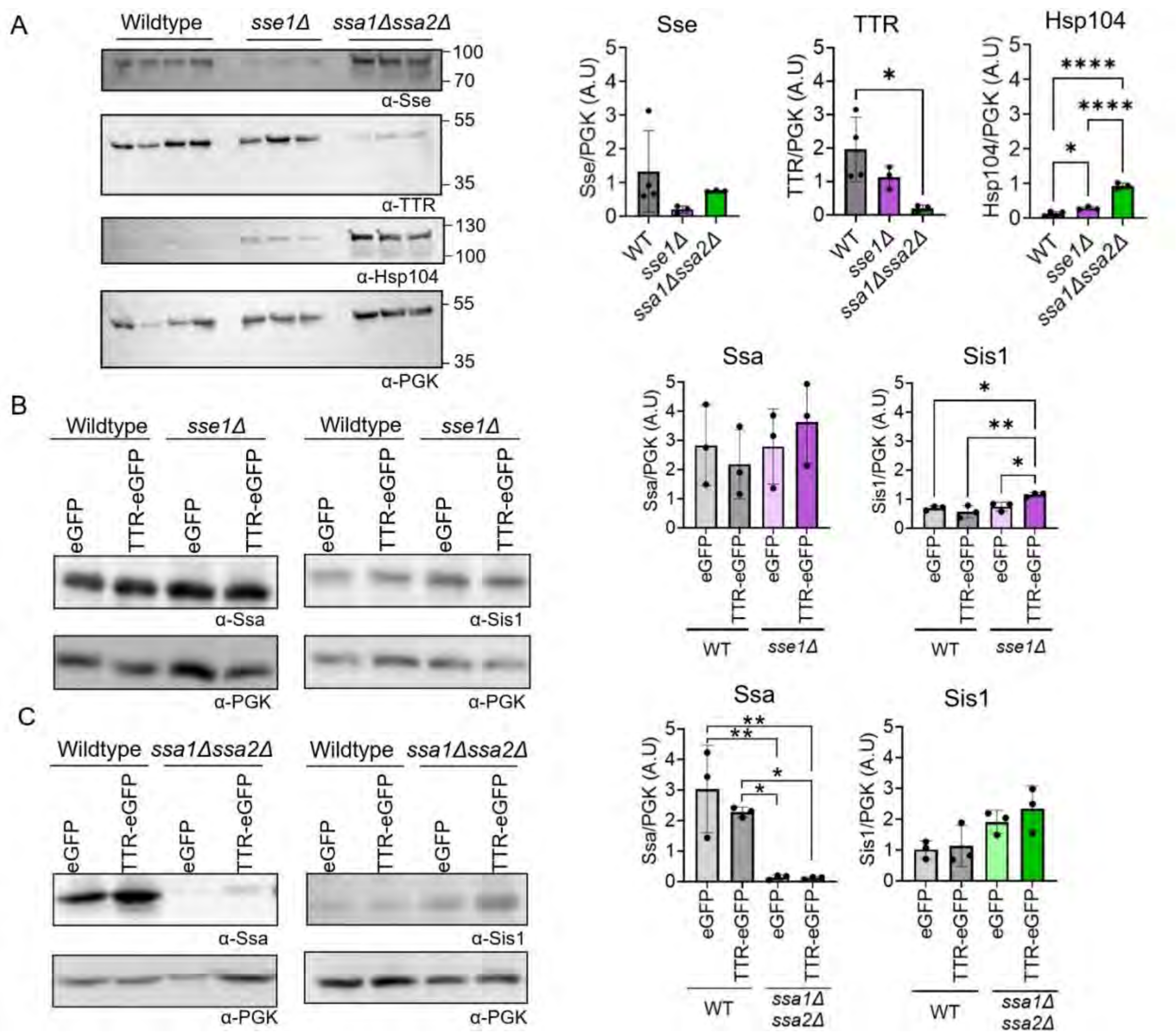


**Supplemental Figure 6. Chaperone and TTR steady state levels in *sse1Δ and ssa1Δssa2Δ* strains*.*** A) Lysates from wildtype (gray bars on graphs), *sse1Δ* (purple bars)*, and ssa1Δssa2Δ* (green bars) strains transformed with TTR-eGFP were analyzed via SDS-PAGE and immunoblotted with monoclonal anti-TTR, anti-Hsp104, anti-Sse (recognizes Sse1 and Sse2) and anti-PGK antibody. B) Wildtype and *sse1Δ* strains subjected to immunoblotting against anti-Ssa (recognizes Ssa1-4), and anti-Sis1 antibodies. C) Same as B except for *ssa1Δssa2Δ* strains. All graphs quantify the steady state levels normalized to PGK. Statistical analyses were performed with a one-way ANOVA and Tukey post hoc test (* $p < 0.05$, ** $p < 0.01$, **** $p < 0.0001$). Unlabeled comparisons are not significant. Dots indicate independent trials, graphed as mean ± SD.

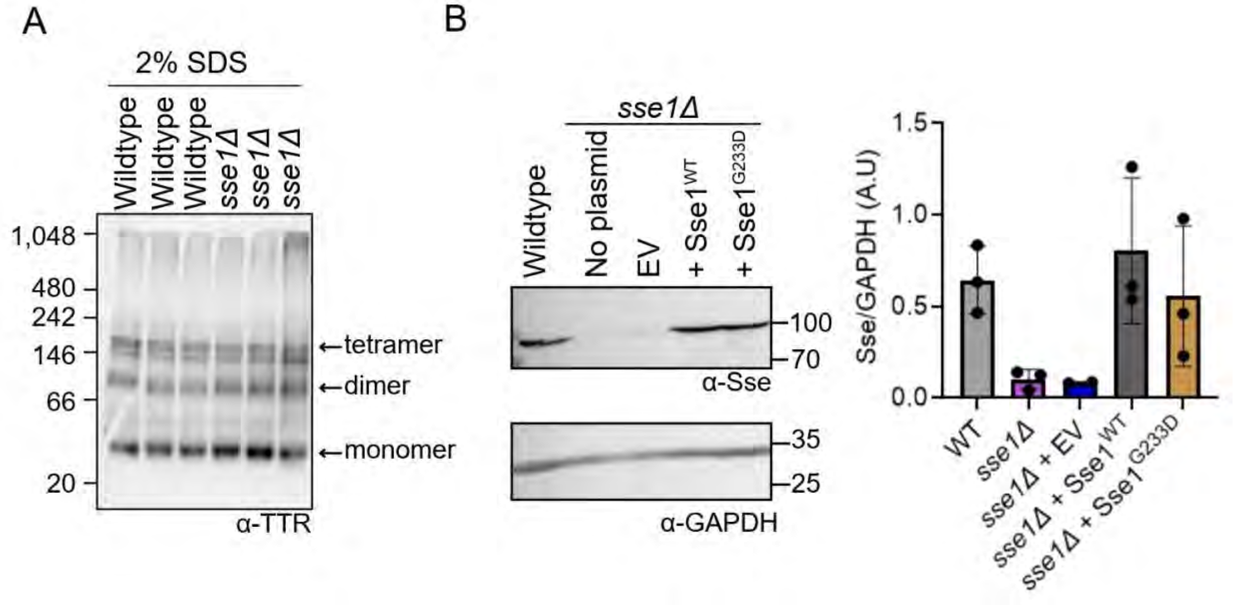


**Supplemental Figure 7. SDS treated *sse1Δ* native PAGE and Sse1 variant steady state levels.** A) Lysates *sse1Δ* strain expressing TTR-eGFP were treated with 2% SDS prior to loading onto 4–20% tris-glycine native PAGE and immunoblotted in polyclonal anti-TTR antibody. Estimated monomeric, dimeric, and tetrameric TTR-eGFP sizes are labeled. B) Lysates of wildtype strains or *sse1Δ* strains expressing either wildtype Sse1 (p3360), Sse1$^{G233D}$ mutant (p3362), or empty vector (p3114) were transformed with TTR-eGFP. Lysates were subjected to Western blot analysis to determine Sse1 steady state levels normalized to GAPDH. Dots indicate independent trials graphed as mean ± SD. Statistical analysis was performed with a one-way ANOVA and Tukey post hoc test (* $p < 0.05$). No comparisons are significant. Dots indicate independent trials graphed as mean ± SD.

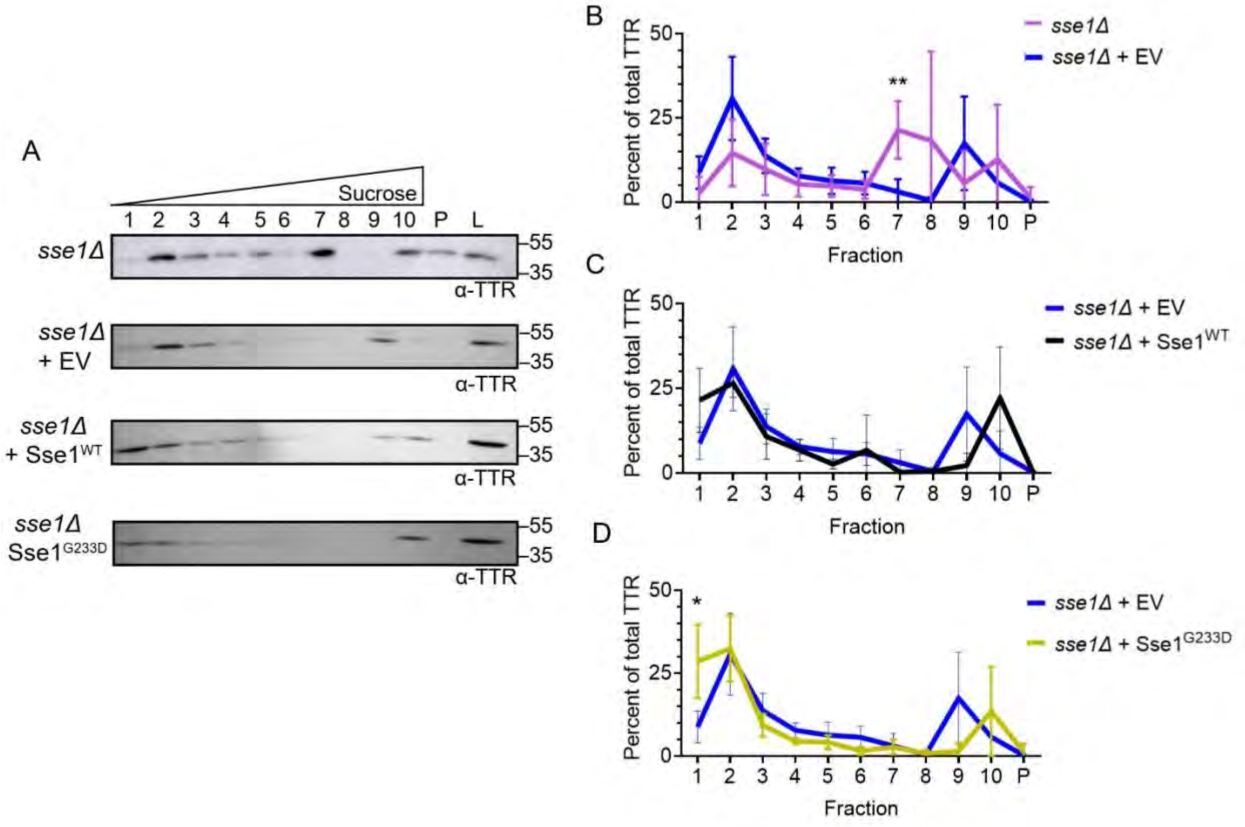


**Supplemental Figure 8. TTR-eGFP is detected in fraction 1 in both *sse1Δ* strains expressing Sse1$^{WT}$ and Sse1$^{G233D}$.** *sse1Δ* strains (M649) were transformed with plasmids expressing either empty vector (p3114), wildtype Sse1 (p3360), or mutant Sse1$^{G233D}$ (p3362) and TTR-eGFP. Sucrose gradient fractionation was performed, and fractions were subjected to Western blot analysis using monoclonal anti-TTR antibody. A) Representative sucrose gradient images of the indicated strains are shown. B) Quantification of the TTR signal from each fraction normalized to the combined TTR signal in fractions 1-10 and pellet of *sse1Δ* (n=6) vs. *sse1Δ* + empty vector (EV; n=5). The mean ± SD is graphed. Statistical analyses were performed with a two-way repeated measures ANOVA, Geisser-Greenhouse correction, and a Sidak multiple comparisons test (* $p < 0.05$). C) Same as B, except *sse1Δ* + EV to *sse1Δ* + Sse1$^{WT}$ (n=4), and D) *sse1Δ* + EV to *sse1Δ* + Sse1$^{G233D}$ (n=4). Unlabeled comparisons are not significant.

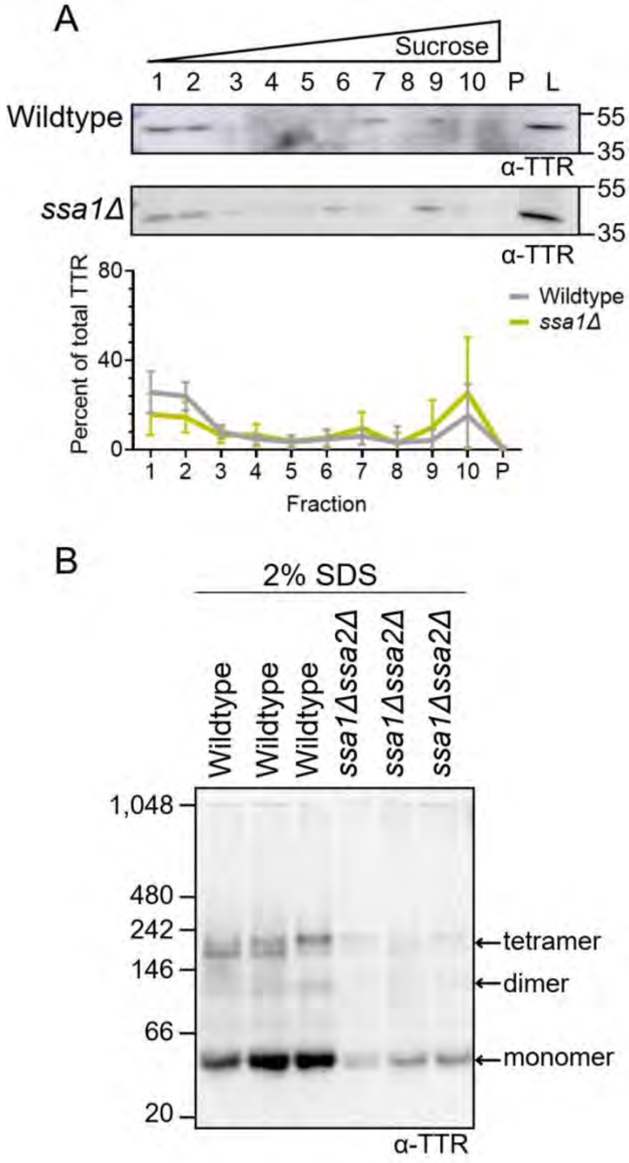


**Supplemental Figure 9. *ssa1Δ* shows no change in TTR-eGFP sedimentation and SDS treatment shows no change in TTR-eGFP in *ssa1Δssa2Δ* native PAGE.** A) Lysates from *ssa1Δ* strains (M652) expressing TTR-eGFP were subjected to discontinuous sucrose gradient centrifugation (10%, 40%, and 60%). Fractions were analyzed by SDS-PAGE and immunoblotted with a monoclonal anti-TTR antibody. Shown is a representative image of five independent trials. The percent of TTR signal normalized to the combined TTR signal in all fractions was quantified and graphed as mean ± SD. Statistical analyses were performed with a two-way repeated measures ANOVA, Geisser-Greenhouse correction, and a Sidak multiple comparisons test (* $p < 0.05$). No comparisons are significant. B) Lysates from *ssa1Δssa2Δ* strains expressing TTR-eGFP were treated with 2% SDS prior to 4–20% tris-glycine native PAGE and immunoblotted with polyclonal anti-TTR antibody. Estimated monomeric, dimeric, and tetrameric TTR-eGFP sizes are labeled.

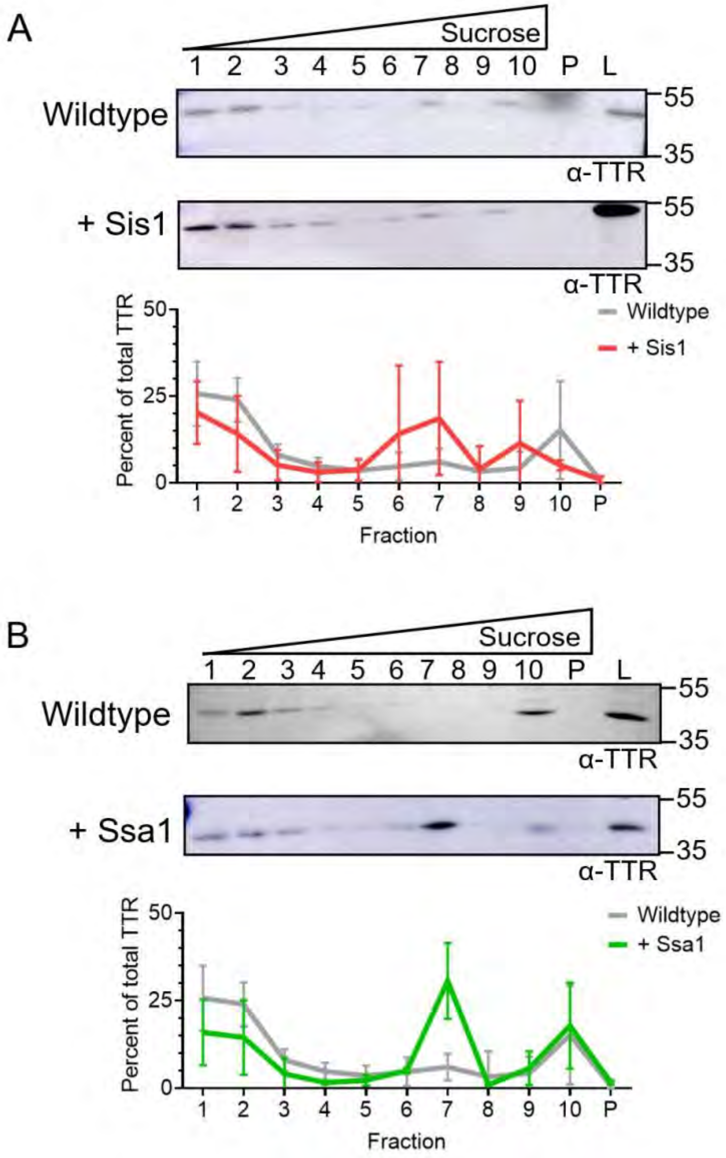


**Supplemental Figure 10. No significant changes detected in TTR-eGFP sedimentation upon Sis1 and Ssa1 overexpression.** Wildtype strains transformed with TTR-eGFP and A) GPD-Sis1 (p3347) or B) GPD-Ssa1 (p3302) were subjected to discontinuous sucrose gradient sedimentation. Fractions were subjected to Western blot analysis using a monoclonal anti-TTR antibody. Shown are representative blots from three independent trials each. The TTR signal from each fraction was normalized to the combined TTR signal in fractions 1-10 and pellet, and the mean ± SD is graphed. Statistical analyses were performed with a two-way repeated measures ANOVA, Geisser-Greenhouse correction, and a Sidak multiple comparisons test (* $p < 0.05$). No comparisons are significant.

**Supplementary Table 1. Yeast strains used in this study**

| Genetic background | Strain number | Genotype | Name in manuscript | Reference |
|---|---|---|---|---|
| 74-D694 | D230 | *MAT**a** ade1-14 ura3-52 leu2-3,112 trp1-289 his3-200 [psi⁻][pin⁻]* | Wildtype | (Chernoff, Lindquist et al. 1995) |
| 74-D694 | M649 | *MAT**a** ade1-14 ura3-52 leu2-3,112 trp1-289 his3-200 sse1::HIS3 [psi-] [pin-]* | *sse1Δ* | This study |
| 74-D694 | M650 | *MAT**a** ade1-14 ura3-52 leu2-3,112 trp1-289 his3-200 sse2::HIS3 [psi-] [pin-]* | *sse2Δ* | This study |
| 74-D694 | M652 | *MAT**a** ade1-14 ura3-52 leu2-3,112 trp1-289 his3-200 ssa1::HIS3 [psi-] [pin-]* | *ssa1Δ* | This study |
| 74-D694 | M674 | *MAT**a** ade1-14 ura3-52 leu2-3,112 trp1-289 his3-200 ssa1::HIS3 ssa2::NatR [psi-] [pin-]* | *ssa1Δ ssa2Δ* | (Buchholz, Martin et al. 2025) |
| 74-D694 | M677 | *MAT**a** ade1-14, ura3-52, leu2-3,112, trp1-289, his3-200, Δsis1::LEU2, TetR-Sis1 74D-694 (TRP plasmid) [pin-]* | *sis1-Δ* [*TETrSIS1*] | (Hines, Higurashi et al. 2011) |

**Supplemental Table 2. Plasmids Used in this Study:**

| Plasmid number | Plasmid name | Yeast Marker | Name in manuscript | Reference |
|---|---|---|---|---|
| p3141 | pAG426GPD-ccdB-EGFP | URA3 (2 micron) | EV-GFP | Addgene plasmid #14204 deposited by Susan Lindquist |
| p3146 | pAG426GPD-TTR-WT-EGFP | URA3 (2 micron) | TTR-GFP | (Knier, Davis et al. 2022) |
| p3299 | pAG425GPD-ccdB | LEU2 (2 micron) | Empty vector for untagged TTR | Addgene plasmid #14154; deposited by Susan Lindquist |
| p3301 | pAG425GPD-TTR | LEU2 (2 micron) | Untagged TTR$^{WT}$ | This study |
| p3235 | pAG426GPD-TTR-WT | URA3 (2 micron) | Untagged TTR$^{WT}$ | This study |
| p3240 | pAG426GPD-TTR-V30M | URA3 (2 micron) | Untagged TTR$^{V30M}$ | This study |
| p3242 | pAG426GPD-TTR-L55P | URA3 (2 micron) | Untagged TTR$^{L55P}$ | This study |
| p3244 | pAG426GPD-TTR-L110Q | URA3 (2 micron) | Untagged TTR$^{L110Q}$ | This study |
| p3302 | pAG415-GPD-Ssa1 | Leu2, (CEN) | GPD-Ssa1 | This study |
| p3114 | pRS314 | TRP1 (CEN) | Empty vector for pRS314 Sse1$^{WT}$ and Sse1$^{G233D}$ | (Sikorski and Hieter 1989) |
| p3360 | pRS314-Sse1$^{WT}$ | TRP1 (CEN) | \Sse1$^{WT}$ | This study |
| p3362 | pRS314-Sse1$^{G233D}$ | TRP1 (CEN) | Sse1$^{G223D}$ | This study |

**Supplementary Table 3 – Antibodies used in this study**

| Antibody | Dilution | Clonality | Vendor | Identifier |
|---|---|---|---|---|
| Pre-albumin | 1:1000 | Monoclonal | Santa Cruz Biotechnology | Cat # sc-377517 |
| TTR 1-147 | 1:1000 | Polyclonal | Invitrogen | Cat # PA5-27220 |
| Hsp104 | 1:2000 | Polyclonal | Enzo Life Sciences | Prod. No. ADI-SPA-1040 |
| Phosphoglycerate Kinase | 1:1000 | Monoclonal | Novex by Life Technologies | Cat # 459250 |
| GAPDH | 1:1000 | Monoclonal | Invitrogen | Cat # MA5-15738 |
| Sis1 | 1:10,000 | Polyclonal | Craig Lab | #66932 |
| Ssa1-4 | 1:10,000 | Polyclonal | Craig Lab | #1173 |
| Green Fluorescent Protein | 1:5000 | Monoclonal | Sigma | Cat # G1546 |
| Anti-Mouse, AP | 1:10,000 | N/A | Sigma Life Sciences | SKU A3562-.5ML |
| Anti-Mouse, HRP | 1:10,000 | N/A | Sigma Life Sciences | SKU A9044-2ML |
| Anti-Rabbit, HRP | 1:10,000 | N/A | Sigma Life Sciences | SKU A9169-2ML |

**Supplementary Table 4. Primers used in this study**

| Generated | ID | Name | Sequence (5'-3') |
|---|---|---|---|
| *sse1Δ* | AM532 | AM532_Sse1Δ sense primer | CCATAAGCAAAAAGTACATTGACA AACAACATTTCTTTAAAAGATGACA GAGCAGAAAGCCCTAGTAAAGC |
| *sse1Δ* | AM533 | AM533_Sse1Δ antisense primer | CGGAAAAAACAATAAAGATCCTTTT CTAGTTACTTTGCTGCATTAACACT ACATAAGAACACCTTTGGTGG |
| *sse1Δ* | AM547 | AM547_Sse1diagnostic 4deletion | CCATTTTAAACTCCCTCTGTC |
| *sse2Δ* | AM558 | AM558_Sse2_sense_deletion | TTTTTTACCTGTAACAGACGTAACC AAAGGATATAATATAATGACAGAGC AGAAAGCCCTAGTAAAGC |
| *sse2Δ* | AM559 | AM559_Sse2_antisense_deletion | AGAATAAAGAGGGAACAATCCAAA TAGACAAAAATTCCGACTACATAAG AACACCTTTGGTGG |
| *sse2Δ* | AM561 | AM561_Sse2_sense_diagnostic4deletion | GCCGTTTAGAGATTTTTATTTATG |
| *ssa1Δ* | AM553 | AM553_Ssa1_sense_deletion | GTATTACAAGAAACAAAAATTCAAG TAAATAACAGATAATATGACAGAGC AGAAAGCCCTAGTAAAGC |
| *ssa1Δ* | AM554 | AM554_Ssa1_antisense_deletion | GACATTTTCGTTATTATCAATTGCC GCACCAATTGGCCTACATAAGAAC ACCTTTGGTGG |
| *ssa1Δ* | AM560 | AM560_Ssa1_sense_diagnostic4deletion | CGTTTCCCAATTCTTACTTAAG |
| *ssa1Δssa2Δ* | AM584 | AM584_Ssa2_NatRdisruption_sense | CCAACAGATCAAGCAGATTTTATAC AGAAATATTTATACAATGGGTACCA CTCTTGACG |
| *ssa1Δssa2Δ* | AM585 | AM585_Ssa2_NatRdisruption_antisense | AGTAAAACTTTTCGGATATTTTACA GGGCGATCGCTAAGCTTAGGGGC AGGGCATGCTC |
| *ssa1Δssa2Δ* | AM220 | NATr AS1 | AAGACGGTGTCGGTGGTGAAGG |
| *ssa1Δssa2Δ* | AM422 | AM422_Ssa2_fwd_at -269 | CCGAGAAGTTCTTCCGATTCAC |
| Sse1$^{WT}$ | AM432 | AM432_Sse1_fwd_at -747 | ACGGTAGGAGATTTGCCACTG |
| Sse1$^{WT}$ | AM433 | AM433_Sse1_rev_200_bp_3'_of_stop | GGGTTTAGGAGACTAATGCGTG |
| Sse1$^{G233D}$ | AM625 | AM625_Sse1_G233D_Mut_Sense | CAAGCATTTTGATGGTAGAGACTT C |
| Sse1$^{G233D}$ | AM626 | AM626_Sse1_G233D_Mut_Anti | TCGCAGGCAGTTCCT |

**References:**


1. Chernoff, Y. O., S. L. Lindquist, B. Ono, S. G. Inge-Vechtomov and S. W. Liebman (1995). "Role of the chaperone protein Hsp104 in propagation of the yeast prion-like factor [psi+]." Science **268**(5212): 880–884.
2. Buchholz, H. E., S. A. Martin, J. E. Dorweiler, C. M. Radtke, A. S. Knier, N. B. Beans and A. L. Manogaran (2025). "Hsp70 chaperones, Ssa1 and Ssa2, limit poly(A) binding protein aggregation." Mol Biol Cell **36**(6): ar66.
3. Hines, J. K., T. Higurashi, M. Srinivasan and E. A. Craig (2011). "Influence of prion variant and yeast strain variation on prion-molecular chaperone requirements." Prion **5**(4): 238-244.
4. Sikorski, R. S. and P. Hieter (1989). "A system of shuttle vectors and yeast host strains designed for efficient manipulation of DNA in Saccharomyces cerevisiae." Genetics **122**(1): 19-27.
